# Macrophages restrict tumor permissiveness to immune infiltration by controlling local collagen topography through a Tcf4-Collagen3 fibrotic axis

**DOI:** 10.1101/2025.01.17.633527

**Authors:** Zoé Fusilier, Franck Simon, Isabel Calvente, Lou Crestey, Alexandra Clément, Mathilde Mathieu, Roude Jean-Marie, Florence Piastra-Facon, Jeyani George Clément, Enola Lumineau, Mattia Tonani, Valeria Manriquez, Livia Lacerda, Perrine de Villemagne, Eliane Piaggio, Vincent Semetey, Sylvie Coscoy, Emanuele Martini, Giorgio Scita, Jean-Christophe Gelly, Johanna Ivaska, Hervé Isambert, Christel Goudot, Paolo Pierobon, Ana-Maria Lennon-Duménil, Hélène D. Moreau

## Abstract

During tumorigenesis, the extracellular matrix (ECM), which constitutes the structural scaffold of tissues, is profoundly remodeled. While the impact of such remodeling on tumor growth and invasion has been extensively investigated, much less is known on the consequences of ECM remodeling on tumor infiltration by immune cells. By combining tissue imaging and machine-learning, we here show that the localization of T lymphocytes and neutrophils, which orchestrate antitumor immune responses, can be predicted by defined topographical features of fibrillar collagen networks. We further show that these collagen topographies result from the activation of a fibrotic pathway controlled by the transcription factor Tcf4 upon depletion of tumor-associated macrophages at late tumor stages. This pathway promotes the deposition of collagen 3 by both tumor and stromal cells, resulting in intermingled collagen networks that favor intra-tumoral T cell and neutrophil localization. Importantly, analysis of human colorectal cancer public bulk RNAseq databases showed a strong correlation between *Tcf4* and *collagen 3*, as well as between the expression of these genes and tumor infiltration by T lymphocytes and neutrophils, attesting the clinical relevance of our findings. This study highlights the key structural role of macrophages on the tumor extracellular matrix and identifies collagen network topographies as a major regulator of tumor infiltration by immune cells.

## Introduction

Tumor progression is dictated by the tug-of-war between cancer and immune cells, and tilting the game toward the immune players has been successfully exploited in different immunotherapeutic strategies^1^. Nevertheless, cancer remains the second leading cause of death worldwide^2^, and immunotherapy successes are still limited to few cancer indications, with low success rates, in particular for solid tumors^3^. Among immune players, the pivotal role of T cells is highlighted by their predictive power on prognosis^4,5^ and response to immunotherapies^6,7^. Indeed, a central mechanism of anti-tumor immune responses is T cell-mediated cytotoxicity^8,9^. As T cells need to reach the cancer cells to exert their function, their frequent exclusion from the tumor core promotes tumor escape from immune-mediated destruction^10,11^. Deepening our understanding of the mechanisms underlying tumor-T cell infiltration is therefore of the upmost importance to design and improve therapeutic strategies.

One element of the tumor microenvironment that has been incriminated in mediating T cell exclusion is the dense extracellular matrix (ECM), in particular collagen, which surrounds tumor islets^12^. The fibrillar collagen network is remodeled during tumorigenesis and evolve over time, leading to distinct topographical organizations, named Tumor-Associated Collagen Signatures (or TACS)^13,14^. At advanced stages, many tumors harbor a dense capsula of collagen fibers aligned to the tumor border (TACS-2). T cells tend to accumulate in these structures, limiting their ability to act on cancer cells (in lung tumors^12,15^; ovarian cancer^15^; pancreatic adenocarcinoma^16^; gastric carcinoma^17^). Yet, the pro-tumoral effect of ECM remodeling is not limited to T cell exclusion. It also supports tumor progression directly, by promoting cancer cell proliferation, survival and metastasis^18^. Especially, reorientation of collagen fibers perpendicularly to the tumor edge (TACS-3) has been shown to promote metastasis^13,14,19^, while chaotic reorientation (TACS-6) promotes multidirectional migration of cancer cells^13^. Whether these reorientations could also affect T cell infiltration remains to be established.

Interestingly, recent studies have shed light on the interplay between tumor ECM remodeling and another major immune player, namely macrophages^20–22^. Tumor-associated macrophages promote ECM deposition indirectly by stimulating fibroblasts and directly by producing ECM-associated molecules themselves^20,20–22^, while also contributing to ECM degradation^23–26^. While tumor-associated macrophages (TAMs) were originally identified as anti-tumoral agents^27^, subsequent studies revealed their more ambiguous role^28^. The complexity of the functions attributed to TAMs grows as their diversity is investigated^29^. Not only macrophages can impede the anti-tumor immune response by promoting an immunosuppressive environment (e.g. by reducing T cell cytotoxic activity^28^), they can also diminish the efficacy of immunotherapies, for example by directly interacting with T cells and retaining them at the periphery^30^ or by capturing anti-PD-1 antibodies and reducing their bioavailability for T cells^31^.

Tumor-associated macrophages have also been shown to impact ECM biophysical properties, namely its stiffness^21^ and topography^20^. Based on these observations, we postulated that macrophages could also limit tumor T cell infiltration by controlling the biophysical properties of the ECM. In this study, we combined imaging and transcriptional analysis to test our hypothesis. We demonstrate that depleting macrophages in already established tumors modulates collagen topographies, rendering specific tumor areas more permissive to both T cells and neutrophils, while others only recruit T cells. This process depends on a fibrotic program in both fibroblasts and tumor cells driven by the transcription factor Tcf4, which promotes the expansion of collagen 3-rich disorganized fibrillar networks at the tumor periphery, favoring a TACS-2-like to TACS-6-like transition, favorable to immune cell infiltration. We further observed that the Tcf4-Collagen 3 axis correlates with T cell and neutrophil numbers in human colon adenocarcinoma, confirming the clinical relevance of our findings. Our study brings fundamental knowledge on the behavior of T cells and neutrophils in complex 3D topographies imposed by the ECM and found in tumors. Through the discovery of an alternative fibrotic pathway in tumors driven by Tcf4, which favors immune infiltration, our study challenges the canonical view of tumor-associated fibrosis associated with T cell exclusion and immunosuppression.

## Results

### Macrophage depletion disrupts fibrillar collagen topography in the tumor

To address whether tumor-associated macrophages could control T cell infiltration through ECM remodeling at late stages of tumorigenesis, we chose a mesenchymal model of colon adenocarcinoma (MC38). This model is known to exhibit a high macrophage infiltration^32^, present an aligned collagen capsula reminiscent of TACS-2, and exclude T cells from the tumor core^33^. MC38 cells were injected subcutaneously in the flank of CD64-hDTR mice^34^. CD64-hDTR mice express the human diphteria toxin receptor (DTR) under the control of the CD64 promotor, which is specific of macrophages and monocytes. This model thus provides a tool to deplete macrophages continuously for 5 days (Fig. 1a), on already well-established tumors (starting at day 9, ≃ 200mm^3^, Fig. S1a). Of note, such prolonged depletion was essential as ECM remodeling, which results from the balance between production and degradation of ECM components, is a relatively slow process^35^. As assessed by F4/80 staining on tumor sections, macrophage depletion was efficient from day 10, with a 60% reduction in macrophage (F4/80^+^ cells) density that reached over 80% at day 14 (Fig. 1b,c). GR1^hi^ monocytes were also efficiently depleted in the blood, with a maximum efficacy of 90% at day 10, while GR1^lo^ monocytes counts were non significantly impacted by DT injection (Fig. S1b-d). We observed a slight and transient, although non-significant, increase in neutrophil counts at day 12, which was completely resolved at day 14, ruling out a systemic inflammation caused by macrophage depletion at the time of our subsequent analysis (Fig. S1e). Of note, macrophage depletion at such a late stage of tumor development did not induce any significant change in tumor growth (Fig. 1d, S1f), allowing direct comparison of control (CD64^WT^) and macrophage-depleted (CD64^DTR^) tumors independently of tumor size.

**Figure 1:**
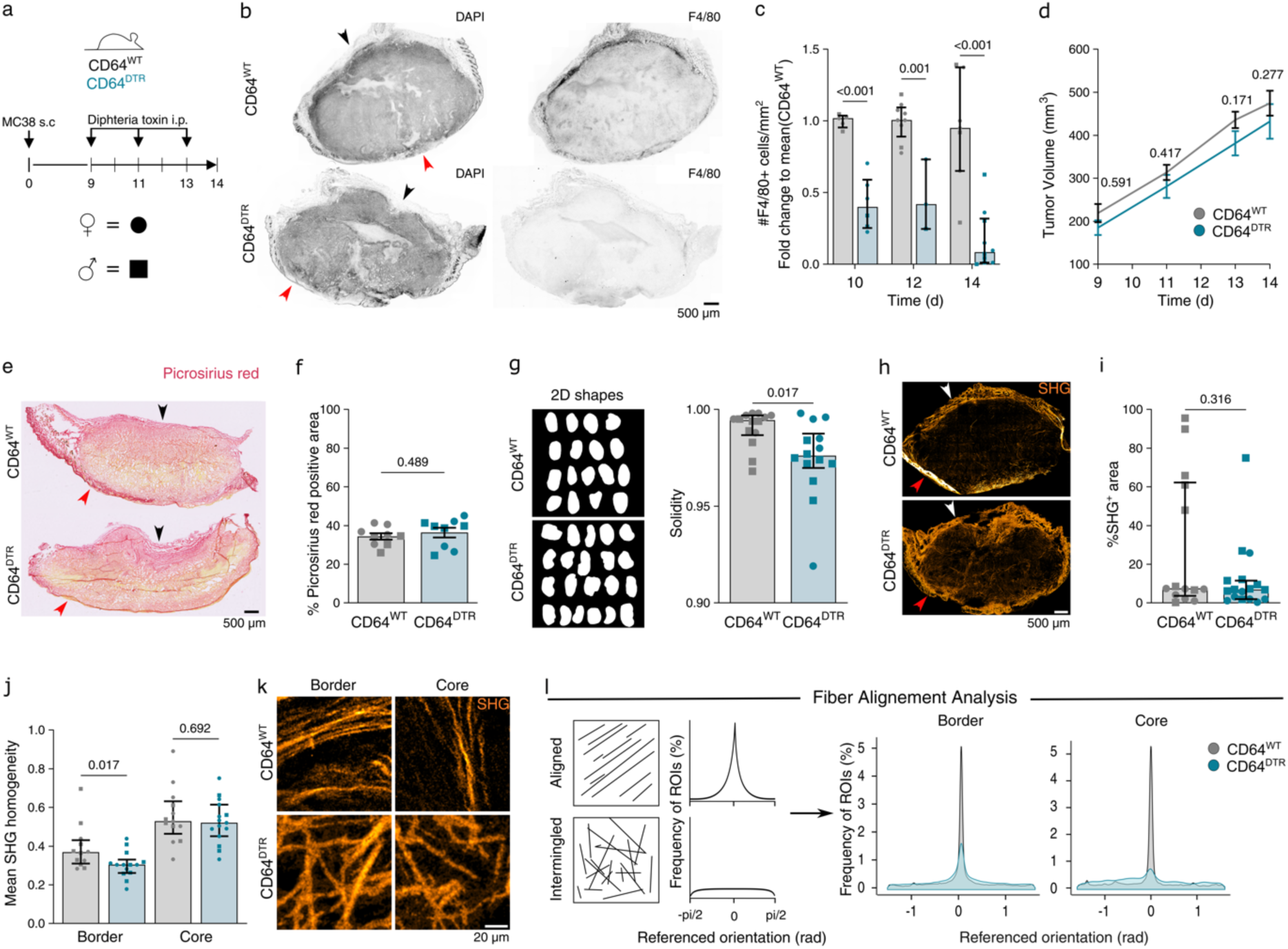
Macrophage depletion disrupts fibrillar collagen topography in the tumor. **(a)** Timeline of macrophage depletion experiments. CDC4^WT^ and CDC4^DTR^ mice were injected intraperitoneally with diphtheria toxin at day S, 11 and 13 after MC38 subcutaneous injection and sacrificed at day 14. Squares represent males and rounds represent females in all graphs. **(b)** Representative epiffuorescence images showing DAPI and F4/80 stainings of CDC4^WT^ (top) and CDC4^DTR^ (bottom) tumors at day 14 (black arrow = tumor border, red arrow = skin). **(c)** Fold change of the number of F4/80^+^ cells per mm^2^ in CDC4^DTR^ tumors over CDC4^WT^ tumors at day 10, 12 and 14 (N=2 independent experiments; median and interquartile range; 2-way-ANOVA test). **(d)** Tumor growth curves from day S to day 14 of CDC4^WT^ and CDC4^DTR^ tumors (N=3 independent experiments, mean and SEM; multiple Mann-Whitney tests). **(e)** Representative brightfield images of picrosirius red staining (black arrow = tumor border, red arrow = skin). **(f)** % of picrosirius red positive (fibrillar collagen) area in CDC4^WT^ and CDC4^DTR^ tumors at day 14 (N=2 independent experiments; median and interquartile range, Mann-Whitney test). **(g)** 2D shapes of tumor slices and associated solidity measurement of CDC4^WT^ and CDC4^DTR^ tumors at day 14 (N=3 independent experiments; median and interquartile range, Mann-Whitney test). **(h)** Representative two-photon images showing SHG signals of CDC4^WT^ and CDC4^DTR^ tumors at day 14 (white arrow = tumor border, red arrow = skin). **(i)** Percentage of SHG^+^ area in CDC4^WT^ and CDC4^DTR^ tumors at day 14 (N=3 independent experiments; median and interquartile range, Mann-Whitney test). **(j)** Mean homogeneity of SHG images at the tumor border and in the core of CDC4^WT^ and CDC4^DTR^ tumors at day 14 (N=3 independent experiments; at least five 320 µm x 320 µm ROIs per tumor, median and interquartile range, Mann-Whitney test). **(k)** Representative two-photon images showing SHG signals at the tumor border and in the core of CDC4^WT^ and CDC4^DTR^ tumors at day 14. **(l)** Analysis schematic (left) and distribution of C3 µm x C3 µm ROIs referenced orientation at the border (0-1000 µm from the border) and in the core (>1000 µm) of the tumor (N=3 independent experiments; C3 µm x C3 µm ROIs; 7707CC ROIs and 10S70S8 ROIS for CDC4^WT^ and CDC4^DTR^ tumors resp.).

Overall collagen density was unaltered by macrophage depletion, as assessed by Picrosirius red staining (Fig. 1e, f). Yet, surprisingly, the macroscopic shape of tumors quantified from tissue sections was distorted: tumor edges were more heterogeneous and often displayed concave shapes in macrophage-depleted tumors, as shown by decreased shape solidity (i.e. ratio of shape area to the area of the smallest enclosing shape, typically a convex hull, Fig. 1g). This was also observed on tumor 3D reconstructions (Fig. S1g). As tissue structure is at least in part dictated by the ECM organization, this prompted us to further characterize the fibrillar collagen architecture by second-harmonic generation (SHG) imaging. Control (CD64^WT^) tumors presented a homogeneous capsula of collagen fibers, all along the edge, on the side opposite to the skin (Fig. 1h, top), while macrophage-depleted (CD64^DTR^) tumors exhibited a more heterogeneous capsula (Fig. 1h, bottom), especially in the concave area. Despite high variability, the percentage of SHG^hi^ regions was unaffected by macrophage depletion (Fig. 1i, consistent with Picrosirius red staining), confirming that overall fibrillar collagen content was not drastically affected by macrophage depletion. To evaluate the homogeneity of fibrillar collagen network, we performed Grey Level Co-occurrence Matrix (GLCM) textural analysis on 350 µm x 350 µm regions of interest (ROIs) of SHG images from both the tumor border and the tumor core. GLCM homogeneity is a measure of the structural uniformity of the collagen fibers within a region, which increases as variation or contrast decreases. This analysis revealed that *(i)* the fibrillar collagen network was more homogeneous in the tumor core as compared to the border, and *(ii)* macrophage depletion decreased fibrillar collagen network homogeneity at the edge of macrophage-depleted tumors (Fig. 1j).

When zooming on collagen fiber organization in control tumors, we observed a thin capsula of aligned collagen fibers parallel to the tumor border (Fig. 1k, top), reminiscent of TACS-2^13^, consistent with the literature^14^. By contrast, the capsula of macrophage-depleted tumors harbored intermingled fibers, with no dominant orientation. This resembles TACS-6, which have been defined as “*chaotically aligned collagen fibers […] without a clear tumor boundary*”^13^ (Fig. 1k, bottom). These observations are consistent with the loss of fiber orientation reported in orthotopic MC38 model in which macrophages have been depleted all along tumorigenesis^20^ as well as in established breast spontaneous tumors^21,22^ where TACS have been originally described^14^. To evaluate quantitatively the degree of collagen fiber alignment, we used a structure tensor analysis to extract the local orientation of collagen fibers^36^, referenced to the mean orientation of a surrounding window of 63 µm x 63 µm (Fig. 1l, left). Structure tensor analysis allows fiber properties analysis without the need of individual segmentation. We observed a decrease in the frequency of ROIs presenting aligned fibers, both in the tumor core and at the edge of macrophage-depleted tumors compared to controls (Fig. 1l, right). In control tumors, both the tumor core and edge exhibited a strong local dominant orientation of collagen fibers, consistent with the homogeneity quantified at a bigger scale by GLCM. Noticeably, macrophage depletion led to a loss of fiber alignment in the tumor core. In addition, at the edge of macrophage-depleted tumors, we observed a mix of aligned and disorganized fibers, consistent with a decreased network homogeneity. Altogether, our observations advocate for a structural role of tumor-associated macrophages in the organization of fibrillar collagen networks: their depletion diversifies local collagen topographies and increases disorganized collagen networks.

### Collagen remodeling is associated to a drastic increase in T cell infiltration in specific network topographies

It has been proposed that the collagen capsula of tumors retains T cells at the edge of tumors, preventing their entry into the tumor core^10,12^. We therefore investigated whether the local changes in collagen organization observed upon macrophage depletion were associated to an increase in tumor permissiveness to T cell infiltration. We found that in our experimental model, T cells (as detected by CD3ε staining, Fig. 2a) infiltrated the tumor only at day 14 (Fig. S2a), consistent with previous reports^33^. This infiltration was increased over two-fold upon macrophage depletion (Fig. 2b), leading to an increased tumor coverage by T cells (Fig. S2b). T cells also penetrated deeper in the tumor, as depicted by their increased proportion in the tumor core (distance to border > 1000 µm, Fig. 2c). To characterize more precisely how infiltrating T cells distribute in the tumor, we defined T^rich^ ROIs as regions with T cell numbers superior to the mean of T cell number in regions with at least one T cell (T^+^). Interestingly, T cells tend to cluster more together in macrophage depleted tumors (increased T^rich^/T^+^ ratio), pointing to local accumulation signals (Fig. 2d). This analysis further revealed that T cells preferentially accumulated in collagen-rich (SHG density > mean of SHG^+^ areas) regions (>90%, Fig. 2e), suggesting that the presence of collagen fibers favors T cell local infiltration and clusterization. Because tissue stiffness is known to be determined by collagen density and topography^37,38^ we further characterized T cell-enriched areas by measuring local stiffness by Atomic Force Microscopy. This analysis highlighted that T cell-enriched regions were stiffer than T cell poor regions, with a markedly stronger difference in macrophage-depleted tumors (x10 in CD64^DTR^ vs x2.3 in CD64^WT^, Fig. S2c). Together, our results show that macrophage depletion leads to a local change in collagen network topography, which is associated to enhanced infiltration by T cells.

**Figure 2:**
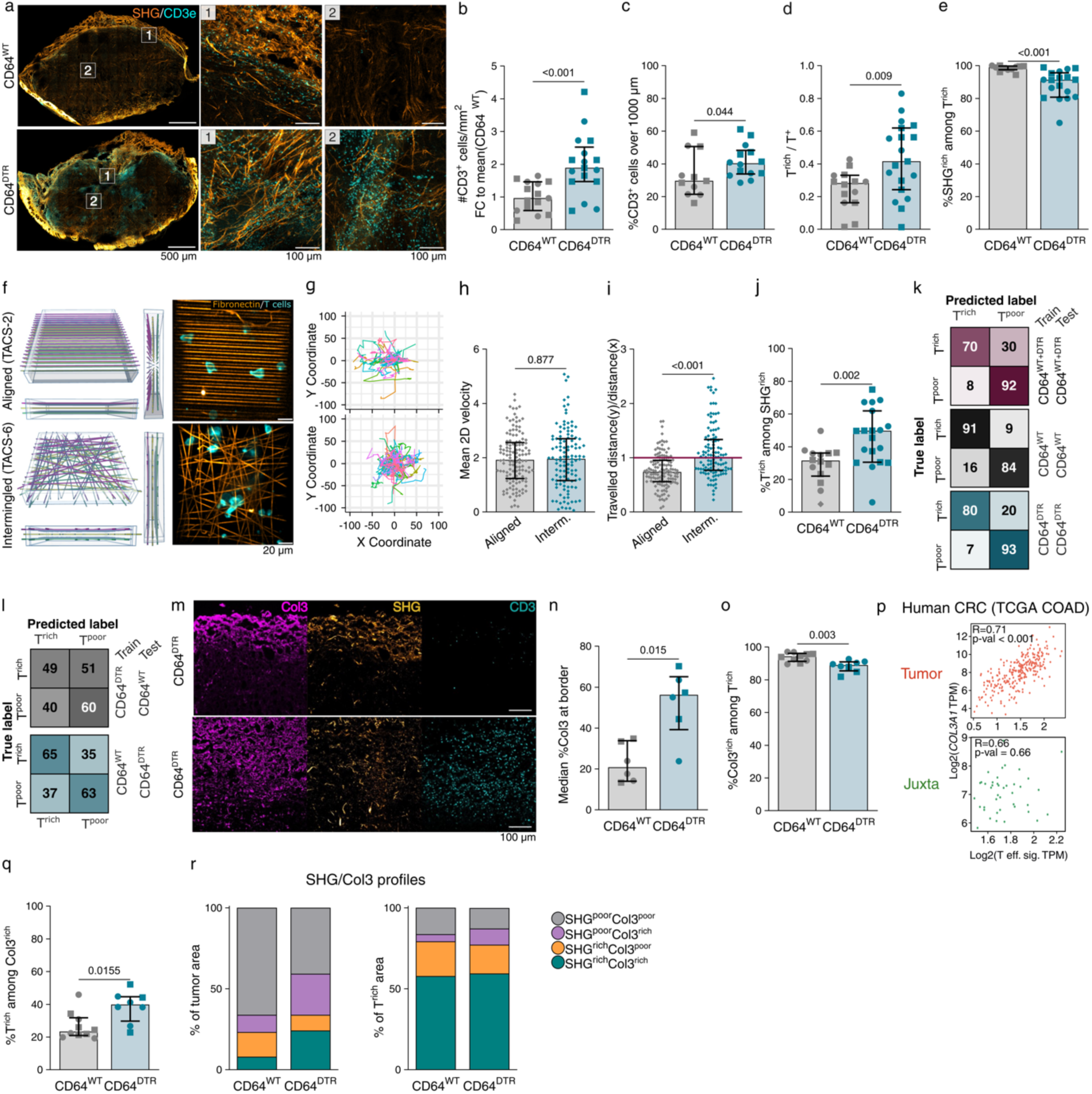
Collagen remodeling is associated to a drastic increase in T cell infiltration in specific network topographies. **(a)** Representative two-photon images showing SHG and CD3ɛ staining at the border (1) and in the core (2) of CDC4^WT^ and CDC4^DTR^ tumors at day 14. **(b)** Fold change of the number of CD3ɛ^+^ cells per mm^2^ in CDC4^DTR^ tumors over CDC4^WT^ tumors at day 14 (N=3 independent experiments; median and interquartile range, Mann-Whitney test). **(c)** %T cells over 1000 µm from tumor border in CDC4^WT^ and CDC4^DTR^ tumors at day 14 (N=3 independent experiments; median and interquartile, Mann-Whitney test). **(d)** Ratio of T^rich^ over T^+^ regions (100 µm x 100 µm tiles) in CDC4^WT^ and CDC4^DTR^ tumors at day 14 (N=3 independent experiments; median and interquartile range, Mann-Whitney test,). **(e)** %T^rich^ regions showing enrichment in SHG signal in CDC4^WT^ and CDC4^DTR^ tumors at day 14 (N=3 independent experiments; median and interquartile range, Mann-Whitney test). **(f)** Design of 3D aligned (TACS-2-like) or intermingled (TACS-C-like) PEG-DA fibers constructions (left). Representative confocal images of ex vivo activated T cells seeded on aligned or intermingled PEG-DA fibers coated with fibronectin (right). **(g)** Representative 1h 2D tracks of T cells seeded in either aligned (top) or intermingled (bottom) structures. **(h)** Mean T cell 2D velocity in 1h in either aligned or intermingled structures (N=3 independent experiments; median and interquartile range, each dot represents the average of instantaneous speed for a single cell, Mann-Whitney test). **(i)** Ratio of maximum distance traveled by T cells in 1h in y over x axis in either aligned (top) or intermingled (bottom) structures (N=3 independent experiments; median and interquartile range, each dot represents a single cell, Mann-Whitney test). **(j)** %SHG^rich^ regions (100 µm x 100 µm tiles) showing enrichment in T cells (N=3; independent experiments; median and interquartile range, Mann-Whitney test). **(k)** Confusion matrices (%) obtained after training and testing random forest classification on a CDC4^WT^ and CDC4^DTR^ merged dataset, and on distinct CDC4^WT^ or CDC4^DTR^ datasets, **(l)** and cross-trained and tested between CDC4^WT^ or CDC4^DTR^ datasets (N=10 models). **(m)** Representative two-photon images of Col3a1, SHG and CD3 from CDC4^WT^ and CDC4^DTR^ tumors at day 14. **(n)** Median %Col3a1 in 100 µm x 100 µm tiles located in the first 500 µm from the tumor border in CDC4^WT^ and CDC4^DTR^ tumors at day 14 (N=2 independent experiments; median and interquartile range, Mann-Whitney test). **(o)** %T^rich^ regions (100 µm x 100 µm tiles) showing enrichment in Col3a1 in CDC4^WT^ and CDC4^DTR^ tumors at day 14 (N=2 independent experiments; median and interquartile range, Mann-Whitney test). **(p)** Spearman correlation of Col3a1 expression and effector T cell signatures in TCGA COAD tumors and juxta-tumoral tissues bulk RNA sequencing dataset. **(q)** %Col3a1^rich^ regions (100 µm x 100 µm tiles) showing enrichment in T cells in CDC4^WT^ and CDC4^DTR^ tumors at day 14(N=2 independent experiments; median and interquartile range, Mann-Whitney test). **(r)** Distribution of SHG^rich^Col3a1^rich^, SHG^rich^Col3a1^poor^, SHG^poor^Col3a1^rich^, SHG^poor^Col3a1^poor^ (100 µm x 100 µm tiles) in CDC4^WT^ and CDC4^DTR^ tumors at day 14 (left). Distribution of SHG^rich^Col3a1^rich^, SHG^rich^Col3a1^poor^, SHG^poor^Col3a1^rich^, SHG^poor^Col3a1^poor^ among T^rich^ regions (100 µm x 100 µm tiles) in CDC4^WT^ and CDC4^DTR^ tumors at day 14 (right) (N=2 independent experiments; median and interquartile range, Mann-Whitney test).

These observations prompted us to hypothesize that by modifying collagen topography, macrophage depletion renders the tumor capsula more permissive to T cell infiltration. To test this hypothesis in a simple and controllable experimental system, we fabricated synthetic 3D structures by two-photon polymerization of PolyEthyleneGlycol-DiAcrylate coated with fibronectin to mimic the ECM, presenting either aligned (along the *x* axis) or intermingled fibrils (Fig. 2f), mimicking TACS-2 and TACS-6 respectively. *Ex vivo* activated T cells were added to these nanofabricated matrices and tracked to quantify their migration (Fig. 2g). We observed that T cells migrated with similar 2D speeds on both types of structures (Fig. 2h). Yet, while T cell trajectories on intermingled fibers were globally isotropic in *x* and *y*, trajectories on aligned fibers exhibited a migration anisotropy along the fiber axis (*x*). Such anisotropic migration limited the distance travelled by T cells in the orthogonal direction (*y*), resulting in increased displacement in *y* relative to *x* on intermingled fibers (Fig. 2i). These results show that T cells better explore their environment when migrating on intermingled fiber topographies as compared to aligned ones. They are consistent with our *in vivo* observations showing an increased penetrance (along the “*y”*-tumor axis) of T cells in the core of macrophage-depleted tumors exhibiting a disorganized collagen capsula (Fig. 2c).

Surprisingly, we observed that while almost all T cell-enriched regions were also collagen-rich (Fig. 2e), not all collagen-rich regions were colonized by T cells (Fig. 2j). This applied to both control (CD64^WT^) tumors, which displayed 31% of collagen-rich regions enriched in T cells in average, and to macrophage-depleted (CD64^DTR^) tumors, in which this number reached 51%. This suggested that *(i)* collagen networks might locally exhibit specific traits that could favor T cell infiltration and/or accumulation, and *(ii)* those traits might be enriched in macrophage-depleted tumors. Such traits could either correspond to specific topographical determinants, or to biochemical signals present in the environment such as chemokines or other attracting factors. To assess whether collagen topography itself could dictate T cell local enrichment, we chose to undertake an unbiased approach that encompasses the complexity of local collagen network topography in tumors and its relationship with T cell infiltration. We therefore analyzed CD3ε/SHG images of control and macrophage-depleted tumors, divided in small ROIs that covered the entire tumor (1867864 ROIs covering 14 control and 19 macrophage-depleted tumors). For each SHG^+^ ROI, we extracted T cell numbers as well as a set of topographical parameters using a structure tensor analysis^36^ at different window sizes (Fig. S2d). T^rich^ ROIs were defined as previously, and the other regions were labeled as T^poor^. This first analysis revealed that none of the individual parameters correlated linearly with local T cell numbers in control (CD64^WT^) and macrophage-depleted (CD64^DTR^) tumors (Fig. S2e). Principal component analysis did not show clear segregation either between the T^rich^ and T^poor^ regions, regardless of the presence of macrophages, suggesting that the relationship between T cell enrichment and collagen topographies is non-trivial (Fig. S2f). This result prompted us to develop a Random Forest model to classify T^rich^ and T^poor^ regions, based exclusively on the SHG topographical features (Fig. S2g). The model was trained on 70% of the dataset, then tested on the remaining 30%, and this process was iterated 10 times on different randomly chosen subsets for cross-validation. To test whether collagen topography was predictive of T cell local enrichment, we performed the random forest classification on the merged dataset of both CD64^WT^ and CD64^DTR^ images. This led to an accurate prediction of T^poor^ regions (92%), and satisfactory prediction of T^rich^ regions (70%) (Fig. 2k top, S2h), demonstrating that association between T cells and specific collagen network features was not random. Yet, we could hope for better performances of the prediction of T^rich^ regions. We reasoned that the topographical differences in collagen networks observed in control and macrophage-depleted tumors (Fig. 1) and/or a different behavior of T cells could confuse the training of the model and thus dampen its predictive power on the merged datasets. To test this hypothesis, we trained and tested separately the model on either CD64^WT^ or CD64^DTR^ images. Strikingly, this increased to 91% and 80% the performances of T^rich^ region prediction in control and macrophage-depleted tumors respectively (Fig. 2k bottom, S2h), suggesting that T cell enrichment have distinct topographical determinants in CD64^WT^ and CD64^DTR^ tumors. To confirm this finding, we cross-trained the model on one dataset (CD64^WT^ or CD64^DTR^) and tested it on the other (CD64^DTR^ or CD64^WT^ resp.). This strategy drastically withdrew the predictive power of the model and led to close-to-random classification (Fig. 2l). These results indicate that the absence of macrophages leads to important changes in global and local collagen topographical determinants and that these collagen traits are sufficient to predict T cell enrichment within tumors. To identify the ECM parameters that are the most likely to influence the selective localization of T cells on the collagen network, we determined the most relevant features of the model using a permutation analysis. We observed that in both CD64^WT^ or CD64^DTR^ images, the presence and the heterogeneity of fibers in terms of shape and intensity (energy), as well as their mutual organization (coherency, angle, and referenced orientation), were the main contributing features for efficient learning and accurate prediction (Fig. S2i), with a stronger contribution of organization in macrophage-depleted (CD64^DTR^) tumors. Hence, the performance of our random-forest classifier model points to an essential role of fibrillar collagen organization in dictating areas of T cell local accumulation, which is modified upon macrophage depletion.

We next investigated what could account for the different organization of collagen fibers in control and macrophage-depleted tumors. Interestingly, collagen 3 (Col3) has been previously associated in tumors with altered fibrillar collagen alignment detected by SHG imaging and dimmer SHG intensity^39–41^. This was reminiscent of our observations in macrophage-depleted tumors, presenting more heterogeneous and intermingled networks as compared to controls. We therefore postulated that local Col3 deposition could account for the increase of T cell-favorable fibrillar collagen networks observed in macrophage-depleted tumors. Consistent with this hypothesis, we found that Col3 deposition was increased in macrophage-depleted (CD64^DTR^) tumors at the tumor border (Fig. 2m, n), while SHG signal of fiber^+^ ROIs was dimmer (Fig. S2j). As hypothesized, T cells localized in Col3^rich^ (> to the mean of Col3^+^ regions) regions (>88%, Fig. 2o). To assess the clinical relevance of our findings, we took advantage of publicly available bulk RNAseq databases of human colorectal tumors from the Cancer Genome Atlas (TGCA, COAD dataset). This analysis confirmed the correlation between effector T cell infiltration signature (Table S1) and COL3A1 expression in tumors (Fig. 2p), consistent with previous work^42^. Of note, this correlation was not observed in juxta-tumoral tissues (Fig. S2p), suggesting tumor-specific cues are required for T cell recruitment.

Yet, similar to what we observed for SHG, all Col3^rich^ regions were not enriched in T cells (Fig. 2q), underlining the contribution of other determinants. We hypothesized that the enrichment of both fibers (SHG^rich^) and collagen 3 may be required. To test this hypothesis, we classified ROIs based on both SHG and collagen 3 content and compared the distribution of the different SHG/Col3 profiles in control and macrophage-depleted tumors (Fig. 2r, left). SHG^poor^ Col3^poor^ regions dominated in both control and macrophage-depleted tumors (70% and 49%, resp.) but were reduced in macrophage-depleted tumors. As predicted, the Col3^rich^ areas (either SHG^rich^ or SHG^poor^) were increased by 2-fold in macrophage-depleted tumors compared to controls. Most interestingly, the distribution of SHG/Col3 profiles of T^rich^ areas (Fig. 2r, right) drastically differed from the SHG/Col3 profiles of tumor areas (Fig. 2r, left), confirming that the distribution of T cells was not random towards fibrillar collagen networks. The SHG^rich^ Col3^rich^ areas now largely dominated (50% and 71% in control and macrophage-depleted tumors resp.), and SHG^rich^ Col3^poor^ followed (23% and 21%). Notably, no significant difference was observed in the SHG/Col3 profiles of regions colonized by T cells between control and macrophage-depleted tumors. This analysis further reinforced and refined the results from our random-forest model: *(i)* T cells accumulate where there are fibers (SHG^rich^), with specific topographies (Col3^rich^), in both control and macrophage-depleted tumors, and *(ii)* the differences in predictions are more likely due to different topographies than a different behavior of the infiltrating T cells.

Our results thus indicate that macrophage depletion promotes a remodeling of fibrillar collagen networks by increasing collagen 3 deposition, leading to an increase in T cell infiltration in specific topographical regions of the tumor.

### Macrophage depletion induces Col3-fibrotic program driven by Tcf4 in cancer cells and fibroblasts

To gain mechanistic insights into how macrophage depletion controls Collagen 3 expression and alters the local topography of fibrillar collagen networks, we performed single-cell RNA sequencing (scRNAseq) of whole control (CD64^WT^) and macrophage-depleted (CD64^DTR^) tumors at day 14, to provide a comprehensive characterization of the tumor microenvironment (Fig. S3a). At low resolution (0.1), we identified four clusters of cells: Non-immune (18716 cells, *Ptprc^-^*), Lymphocytes (1586 cells, *Ptprc^+^, Cd3e^+^*), Monocytes/Macrophages/Dendritic Cells (9777 cells, *Ptprc^+^, Itgam^+^, Fcgr1^+^, H2-Ab1^+^*) and Granulocytes (1826 cells, *Ptprc^+^, Csf3r^+^, S100a8^+^*) (Fig. 3a, S3b, Table S2). Importantly, the analysis of the Mo/Mac/DC cluster confirmed the specific depletion of macrophages, which exhibit a classical tumor-associated signature^43,44^(*Fcgr1^+^ Adgre1^+^ Mrc1^+^ ApoE^+^,* Fig. S3c-f, Table S3).

**Figure 3:**
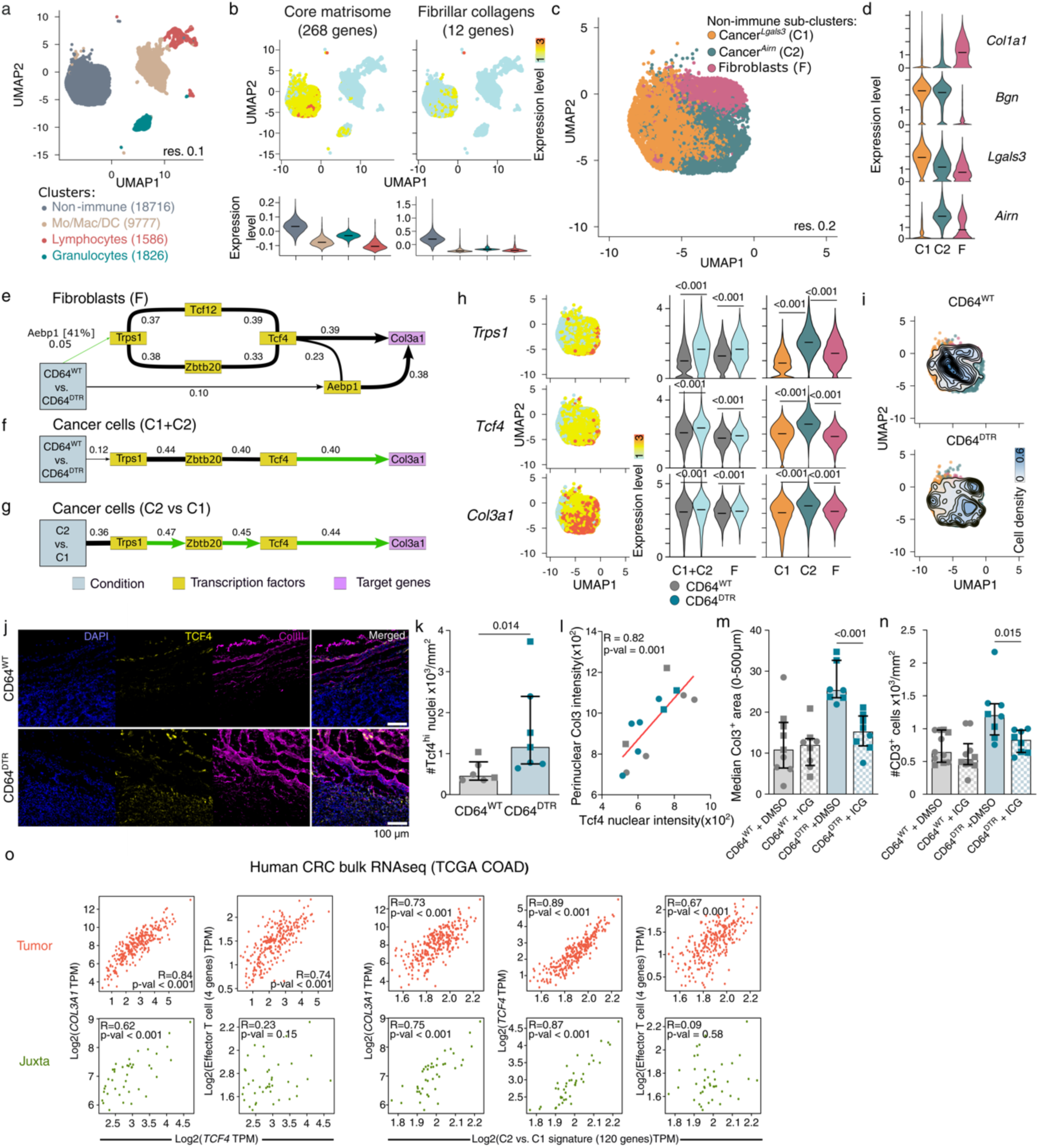
Macrophage depletion induces Col3-fibrotic program driven by Tcf4 in cancer cells and fibroblasts. **(a)** UMAP showing low-resolution clustering of merged scRNAseq data from CDC4^WT^ and CDC4^DTR^ tumors from two independent experiments (For batch #1, live cells (DAPI^-^) were sorted by ffow cytometry; for batch #2, DAPI^-^CD45^-^, DAPI^-^CD45^+^CDC4^+^, DAPI^-^CD45^+^CDC4^-^ populations were sorted by ffow cytometry and sequenced separately). 4 clusters were identified by differential gene expression analysis (DGEA): Non-immune cells, Mo/Mac/DCs, Lymphocytes and Granulocytes. **(b)** UMAPs and violin plots showing core matrisome and fibrillar collagen signature expression levels in low-resolution clusters. **(c)** UMAP showing sub-clustering of the non-immune cluster from CDC4^WT^ and CDC4^DTR^ tumors (batch #1 and batch #2 CD45^-^ samples). DGEA identified 3 clusters: Cancer^Lgals3^ (C1), Cancer^Airn^ (C2), and Fibroblasts (F). **(d)** Expression levels of classical markers used to identify cancer (Bgn, Lgals3, Airn) and fibroblast (Col1a1) clusters. **(e)** Transcriptional network induced by macrophage depletion in fibroblasts and **(f)** in cancer cells (C1+C2) identified by MIIC analysis (Top 50 TFs + 44 collagen genes). **(g)** Transcriptional network induced by cancer cell clustering (C2 vs. C1) independently of macrophage depletion identified by MIIC analysis (top 75 TFs + top 535 genes). **(h)** Expression levels of Trps1, Tcf4 and Col3a1 in non-immune clusters in CDC4^WT^ and CDC4^DTR^ tumors (line at median, Wilcoxon test). **(i)** UMAP of non-immune clusters showing cell densities in CDC4^WT^ and CDC4^DTR^ tumors datasets. **(j)** Representative epiffuorescence of DAPI (blue), Tcf4 (yellow) and Col3a1 (magenta) at the tumor border of CDC4^WT^ and CDC4^DTR^ tumors. **(k)** Number of Tcf4^high^ nuclei in CDC4^WT^ and CDC4^DTR^ tumors (N=2 independent experiments; median and interquartile range, Mann-Whitney test). **(l)** Spearman correlation of mean Tcf4 nucleic intensity and mean Col3a1 perinuclear intensity (5 µm around nucleus). Tcf4 nucleic intensity and Col3a1 cytoplasmic intensity were calculated for each nucleus in the tumor and averaged by tumor (N=2 independent experiments). **(m)** Median %Col3a1 in 100 µm x 100 µm tiles located in the first 500 µm from the tumor border in CDC4^WT^ and CDC4^DTR^ tumors at day 14 treated with Tcf4 inhibitor ICG-001 or DMSO (N=2 independent experiments; median and interquartile range, Mann-Whitney test). **(n)** Number of T cells per mm^2^ in CDC4^WT^ and CDC4^DTR^ tumors at day 14 treated with Tcf4 inhibitor ICG-001 or DMSO (N=2 independent experiments; median and interquartile range, Mann-Whitney test). **(o)** Spearman correlation between expression of TCF4, COL3A1 and effector T cell signature (left) and between Cancer^Airn^ signature (C2 vs. C1) expression and COL3A1, TCF4 and effector T cell signature (right) in TCGA COAD tumors and juxta-tumor tissues bulk RNA sequencing dataset.

We then focused on ECM genes to identify candidates for the regulation of collagen networks upon macrophage depletion. As expected, the Non-immune cell cluster expresses the highest levels of Core matrisome genes (268 genes^45^) and fibrillar collagen (12 genes, GO:0005583^46^) genes (Fig. 3b, Table S4). By sub-clustering of the non-immune cluster at higher resolution (0.2), we identified three different clusters within the Non-immune subset: one cluster of Fibroblasts (F, defined by its high expression of *Col1a1*), and two clusters of cancer cells both expressing the cancer marker *Bgn* and specific gene signatures: Cancer^Lgals3^ (C1, whose most differentially expressed gene was *Lgals3*) and Cancer^Airn^ (C2, whose most differentially expressed gene was *Airn*) (Fig. 3c,d, Table S5). To decipher the pathway leading from macrophage depletion to fibrillar collagen reorganization, we reconstructed the causal gene regulatory network from the scRNAseq data. To do so, we took advantage of the Multivariate Information-based Inductive Causation (MIIC) algorithm that infers direct and possibly causal relations, as well as indirect pathways, in large networks^47,48^. For fibroblasts on one hand, and total cancer cells (C1+C2) on the other, we selected the transcription factors sharing the highest mutual information with the *Condition* (CD64^WT^ or CD64^DTR^), i.e. whose changes in expression are more strongly influenced by macrophage depletion. We then ran the algorithm on the scRNAseq dataset restricted to the *Condition*, the list of most informative transcription factors (top 50 or top 250 TFs), and total collagen genes (44 genes^45^). Strikingly, the simplified regulatory networks obtained for fibroblasts and cancer cells shared many players. More specifically, we found that the *Condition* (macrophage depletion) controls the expression of *Trps1* (as well as *Aebp1* in fibroblasts), which controls the expression of *Tcf4*, that in turn regulates the expression of *Col3a1* and several other collagen genes (Fig. 3e-f, S3g, h, *Full networks available here: <u>Fibroblasts</u>, <u>Cancer</u> <u>cells</u>*). Interestingly, unbiased MIIC analysis on the top 75 most informative TFs between C1 and C2 clusters (independently of macrophage depletion) and including also the top 535 non-TF genes (selected with the same mutual information threshold as TFs) recovered the same regulatory axis (Fig. 3g, *full network available <u>here</u>*). This result suggests that the *Trps1*-*Tcf4*-*Col3a1* transcriptional axis is associated to C2 phenotype, independently of macrophage depletion. Accordingly, gene expression profiles confirmed that macrophage depletion significantly increases *Trps1*, *Tcf4* and *Col3a1* expression in all non-immune clusters (Fig. 3h). Consistent with MIIC analysis, we also found that the C2 cluster was the biggest contributor of *Trps1*, *Tcf4* and *Col3a1* when compared to the other non-immune clusters (Fig. 3h) and overall expressed more core matrisome and fibrillar collagen genes than the C1 cluster (Fig. S3i). The C2 cluster was also enriched in macrophage-depleted (CD64^DTR^) as compared to control tumors, suggesting that macrophage depletion favors a pro-fibrotic/ECM program in cancer cells (Fig. 3i, Fig. S3j). Accordingly, binding motifs enrichment analysis in the promoter regions of the differentially up-regulated genes (Log2FC > 0.25) between CD64^DTR^ and CD64^WT^ in both fibroblasts and cancer cells (C1+C2) showed that they all exhibited a significant enrichment in *Tcf4* binding motif (Fig. S3k, Table S6), identifying Tcf4 as a possible driver of C2 phenotype. This was also observed when comparing C2 over C1, i.e. independently from the *Condition*. Of note, we also found significant enrichment of *Tcf3* and *Tcf12* binding motifs which showed a high degree of similarity with Tcf4 binding motif (Fig. S3k, l). Tcf4, Tcf3 and Tcf12 are bHLH type I TFs involved in the Wnt signaling pathway^49,50^. While Tcf4 binding motif was not directly identified in the *Col3a1* promoter, we found the binding motif of both Tcf12 (also identified by MIIC) and Tcf3. In conclusion, we identified that macrophage depletion induces an alternative fibrotic program driven by *Tcf4* and inducing the up-regulation of *Col3a1* in cancer cells and fibroblasts.

To validate experimentally the regulatory pathway predicted by MIIC, we performed immunofluorescence staining for Tcf4 and Col3. Density of Tcf4^+^ nuclei was increased in macrophage-depleted (CD64^DTR^) tumors compared to control (CD64^WT^) tumors (Fig. 3j-k). Additionally, Tcf4^+^ nuclei spatially associated with perinuclear Col3a1 (Fig. 3l), supporting our mechanistic model. To formally establish the causal link between Tcf4 and Col3, we combined macrophage depletion with the inhibition of Tcf4-mediated transcription (using ICG-001^51^). While Tcf4 inhibition had no effect on tumor growth (Fig. S3m), we observed a decrease in Col3 density at the tumor edge in macrophage-depleted tumors, but not in controls (Fig. 3m), suggesting that this TF might indeed account for increased Col3 in macrophage-depleted tumors. Accordingly, inhibition of Tcf4 also reduced T cell infiltration (Fig. 3n) and tumor coverage by T cells (Fig. S3n) in macrophage-depleted tumors, with no noticeable effect on control tumors, further reinforcing this conclusion.

Analysis of the human colorectal tumor datasets (TGCA, COAD) revealed a significant correlation between the expression of *TCF4* and *COL3A1,* as well as *TCF4* and effector T cell signature (Fig. 3o, left, Table S1). *TCF4*, *COL3A1* and effector T cell signature also correlated with the signature of the C2 cluster (human orthologs of differentially expressed genes between C2 and C1 with Log2FC > 1) (Fig. 3o, right, Table S7). Notably, the expression of *TCF4*, *COL3A1* and the signature of the C2 cluster also correlated in the juxta-tumoral tissues, but not with effector T cell signature, supporting that tumor-specific additional determinants are attracting T cells locally (Fig. 3o).

Altogether, our results support that macrophage depletion at late stage of tumorigenesis drives a Tcf4-dependent fibrotic program in cancer cells and fibroblasts, leading to topographical remodeling of fibrillar collagen networks through the deposition of collagen 3, which favors local T cell accumulation. They further suggest that this pathway may be at play in colorectal human tumors, supporting the clinical relevance of our findings.

### Fibrillar collagen topography dictates tumor overall permissiveness to immune infiltration

We then wondered whether the fibrillar collagen remodeling specifically influenced the recruitment of distinct T cell subsets or exerted a broader effect on lymphocyte populations. To address this, we isolated the lymphocyte cluster identified at low resolution (0.1, Fig. 3a) and performed sub-clustering at higher resolution (0.3). We identified 5 lymphocyte sub-clusters: CD8 exhausted T cells (T CD8ex: *Cd3e^+^, Cd8a^+^, Pdcd1^+^, Lag3^+^, Havcr2^+^, Tox^+^, Ctla4^+^*), a mix of CD8^+^ and CD4^+^ stem-like T cells (T stem-like: *Cd3e^+^, Tcf7^+^, Lef1^+^*), B cells (B: *Cd3e^-^, Cd19^+^*), regulatory T cells (Tregs: *Cd3e^+^, Cd4^+^, Pdcd1^+^, Lag3^+^, Havcr2^+^, Tox^+^, Il2ra^+^, Foxp3^+^, Ctla4^+^, Itgae^+^*), and a cluster of NK cells (NKs: *Cd3e^-^, Klrbc1^+^, Gzma^+^, Gzmb^+^*) (Fig. 4a, b, Table S8). Further focusing on T cells (*CD3e^+^*), we performed differential gene expression analysis using the DESeq2 package^52^ on the 3 T cell clusters we identified (T CD8ex, T stem-like, Tregs), between control (CD64^WT^) and macrophage-depleted (CD64^DTR^) tumors. Surprisingly, none of the CD3^+^ clusters showed a marked difference in terms of transcriptome between control (CD64^WT^) and macrophage-depleted (CD64^DTR^) tumors (Fig. 4c, Table S9; 3, 5 and 2 genes with Log2FC >1 and adjusted p-value < 0.05 for T CD8ex, T stem-like, Tregs resp.). This shows that macrophage depletion does not alter the transcriptional profile of infiltrating T cell subsets. However, macrophage depletion could bias T cell infiltration toward a specific cluster. To address this question, we performed multiplex imaging to quantify the number of different T cell clusters in control and macrophage-depleted tumors: CD8 T cells (CD3^+^, CD8^+^, PD-1^+^), stem-like T cells (CD3^+^, TCF1^+^), Tregs (CD3^+^, CD4^+^, FOXP3^+^), Thelper (CD3^+^, CD4^+^, FOXP3^-^) and cycling T cells (CD3^+^, KI67^+^). Notably, regardless of macrophage depletion, the majority of tumor infiltrating T cells were CD8^+^, which present a terminally exhausted phenotype according to the scRNAseq analysis, consistent with the lack of tumor control. Most importantly, we found that macrophage depletion did not change the proportion of each T cell cluster, but rather increased T cell numbers in all of them (Fig. 4d-f). Of note, this analysis also showed that macrophage depletion did not affect the proportion of cycling Ki67^+^ T cells, and only moderately increased their number, ruling out that preferential local T cell enrichment in macrophage-depleted tumors was the result of local proliferation upon infiltration. These results indicate that tumor permissiveness to T cells is not biased toward a specific cluster but rather affects all T cell subtypes equally. Consistently, *COL3A1* and *TCF4* expression also correlated with multiple T cell subset signatures (exhausted T cells and Th1-like cells for example) in bulk RNAseq of human colorectal tumors (TCGA, COAD) (Fig. 4g, Table S1). Notably, these results uncouple the effect of tumor-associated macrophages on T cell activation and migration. They further suggest that the topographical changes in collagen networks induced by late depletion of macrophages promote pan-T cell infiltration without modifying their phenotype.

**Figure 4:**
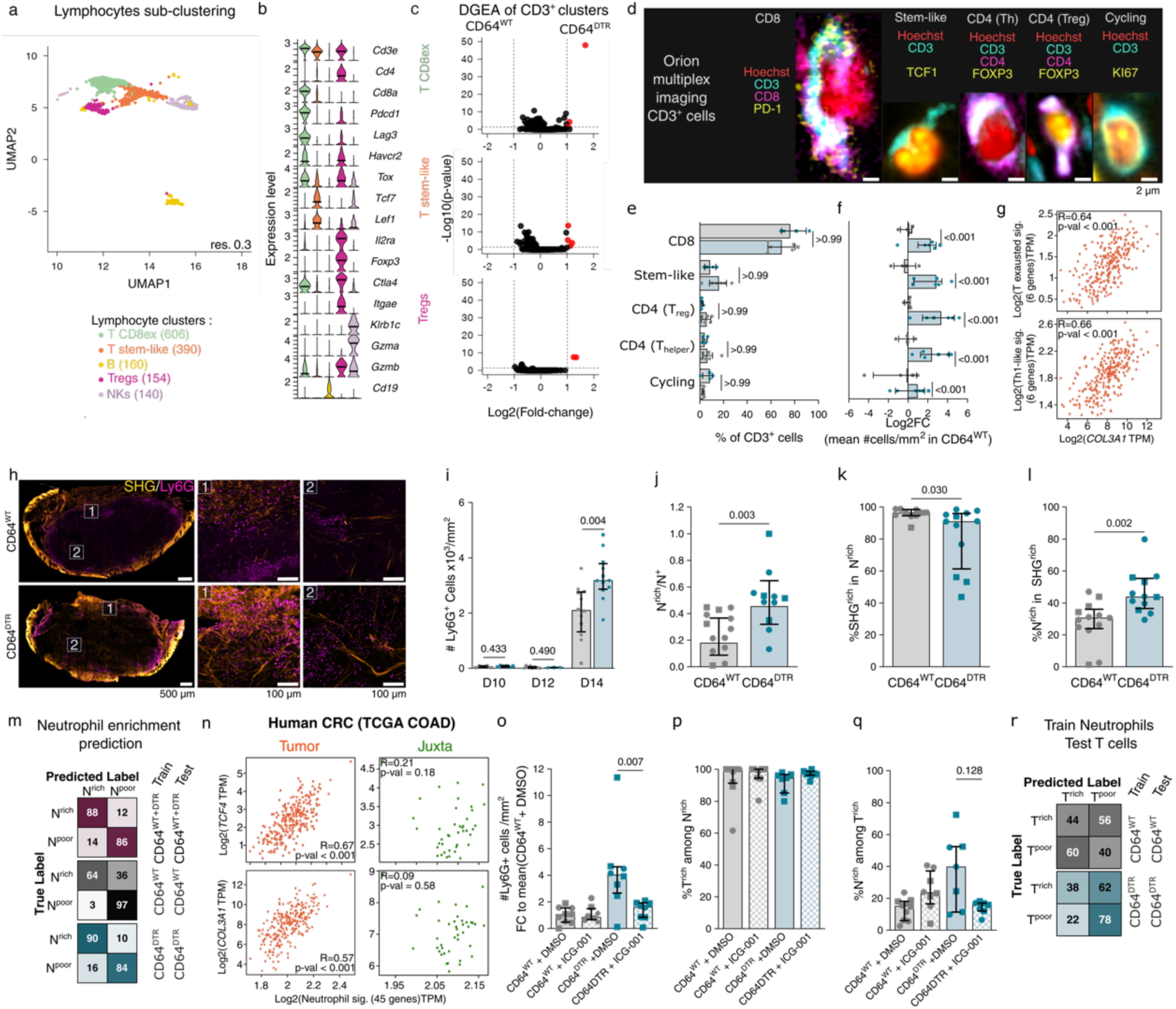
Fibrillar collagen topography dictates tumor overall permissiveness to immune infiltration. **(a)** UMAP showing sub-clustering of lymphocyte cluster from CDC4^WT^ and CDC4^DTR^ tumors (batch #1 and batch #2 DAPI^-^CD45^+^CDC4^-^ samples). 5 clusters were identified by DGEA: T CD8^+^ex, T stem-like, B cells (B), Tregs and NK cells (NK). **(b)** Expression levels of classical T cell markers used to identify lymphocyte clusters. **(c)** DGEA of CD3^+^ clusters CDC4^WT^ and CDC4^DTR^ tumors (black dots show genes with log2FC < 1 and p-adjusted value >0,05, red dots show genes with log2FC > 1 and p-adjusted value <0,05). **(d)** Representative images of Orion multiplex analysis of CDC4^DTR^ and CDC4^WT^ tumors showing examples of CD8, Stem-like, regulatory, helper and cycling T cells. **(e)** Proportions of CD3^+^ clusters/mm^2^ in CDC4^DTR^ and CDC4^WT^ tumors (N=2 independent experiments; median and interquartile range, 2-way-ANOVA test). **(f)** Log2(FC) of number of CD3^+^ clusters/mm^2^ in CDC4^DTR^ and CDC4^WT^ tumors N=2 independent experiments; median and interquartile range, 2-way-ANOVA test). **(g)** Spearman correlation between expression of COL3A1 and exhausted and Th-1-like T cell signatures in TCGA COAD tumors bulk RNA sequencing dataset. **(h)** Representative two-photon images showing SHG and LyCG staining at the border (1) and in the core (2) of CDC4^WT^ and CDC4^DTR^ tumors at day 14. **(i)** Number of LyCG^+^ cells/mm^2^ in CDC4^WT^ and CDC4^DTR^ tumors at day 10,12,14 (N=2 independent experiments for day 10 and 12; N=3 independent experiments for day 14; median and interquartile range, Mann-Whitney test). **(j)** Ratio of neutrophil^rich^ over neutrophil^+^ regions (100 µm x 100 µm tiles) in CDC4^WT^ and CDC4^DTR^ tumors at day 14 (N=3 independent experiments; median and interquartile range, Mann-Whitney test). **(k)** %Neutrophil^rich^ regions (100 µm x 100 µm tiles) showing enrichment in SHG signal in CDC4^WT^ and CDC4^DTR^ tumors at day 14 (N=3 independent experiments; median and interquartile range, Mann-Whitney test). **(l)** %SHG^rich^ regions (100 µm x 100 µm tiles) showing enrichment in neutrophils in CDC4^WT^ and CDC4^DTR^ tumors at day 14 (N=3 independent experiments; median and interquartile range, Mann-Whitney test). **(m)** Confusion matrices (%) obtained after training and testing random forest classification on a CDC4^WT^ and CDC4^DTR^ merged dataset, on distinct CDC4^WT^ or CDC4^DTR^ datasets (N=10 models). **(n)** Spearman correlation between expression of TCF4, COL3A1 and neutrophil signature in TCGA COAD tumors and juxta-tumor tissues bulk RNA sequencing dataset. **(o)** Number of neutrophils per mm^2^ in CDC4^WT^ and CDC4^DTR^ tumors at day 14 treated with Tcf4 inhibitor ICG-001 or DMSO (N=2 independent experiments; median and interquartile range, Mann-Whitney test). **(p)** %Neutrophil^rich^ regions (100 µm x 100 µm tiles) showing enrichment in T cells in CDC4^WT^ and CDC4^DTR^ tumors at day 14 treated with Tcf4 inhibitor ICG-001 or DMSO (N=2 independent experiments; median and interquartile range, Mann-Whitney test). **(q)** %T^rich^ regions (100 µm x 100 µm tiles) showing enrichment in neutrophils in CDC4^WT^ and CDC4^DTR^ tumors at day 14 treated with Tcf4 inhibitor ICG-001 or DMSO (N=2 independent experiments; median and interquartile range, Mann-Whitney test). **(r)** Confusion matrices (%) obtained after training on neutrophil^rich^ topographies and tested on T^rich^ topographies in CDC4^WT^ or CDC4^DTR^ datasets (N=10 models).

We therefore investigated whether the increased tumor permissiveness in macrophage-depleted tumors was restricted to T lymphocytes or whether it also concerned other immune cells. Our scRNAseq revealed an important increase in the infiltration of granulocytes in macrophage-depleted tumors (Fig. S4a), prompting us to assess whether collagen network remodeling induced by macrophage depletion could also impact neutrophil infiltration. We observed a strong infiltration of neutrophils (assessed by Ly6G staining) in control tumors at day 14, which was further enhanced by macrophage depletion (Fig. 4h,i). Similar to our observations for T cells (Fig 2a-e, S2a, b), neutrophils tended to cluster together (Fig. 4j), covered larger areas of the tumor (Fig. S4b) and localized in SHG^hi^ regions (Fig. 4k). Yet, as for T cells, not all SHG^hi^ regions were enriched in neutrophils (Fig. 4l). We therefore postulated that SHG^hi^ regions that were enriched in neutrophils may exhibit specific topographical features leading to their local accumulation, as observed for T cells. To test this hypothesis, we applied our random-forest strategy to test whether fibrillar collagen features assessed by SHG could predict neutrophil localization. Strikingly, a “neutrophil-trained” model on the merged dataset (CD64^WT^ + CD64^DTR^) predicted accurately Neutrophil^rich^ and Neutrophil^poor^ regions (88% and 86% resp., Fig. 4m, top). Training the model separately on CD64^WT^ and CD64^DTR^ led to similar performances, except for the prediction of Neutrophil^rich^ regions by the model trained on CD64^WT^ only (64%), likely due to the smaller size of the dataset (Fig. 4m, bottom). This suggests that the determinants of Neutrophil^rich^ regions are shared between control and macrophage-depleted tumors, and that increased neutrophil recruitment in the latter could be due to an increase in permissive areas.

This result prompted us to investigate whether neutrophil infiltration also relied on the Tcf4-Col3 fibrotic pathway as for T cells, Col3 areas being shared by both control and macrophage-depleted tumors but considerably increased in macrophage-depleted ones (Fig. 2n, r). Consistent with this hypothesis, *COL3A1*, *TCF4*, Cancer^Airn^ (C2) and neutrophil signatures (Table S1) correlated in human tumor COAD TCGA dataset (Fig. 4n, Fig. S4c). As for T cells, neutrophil signature did not correlate with *COL3A1* and *TCF4* in juxta-tumoral tissues, highlighting that tumor specific determinants are required for their infiltration (Fig. 4n). Quantification of neutrophil infiltration after chemical inhibition of Tcf4 with ICG-001resulted in the loss of neutrophil increase in number (Fig. 4o) as well as coverage (Fig. S4d) in macrophage-depleted tumors, similar to what we observed for T cells. Of note, when comparing T cell and neutrophil localization, we observed that all neutrophil-rich areas overlapped with T cell-enriched areas (Fig. 4p), while only a minor fraction of T cell-enriched areas were also enriched in neutrophils (Fig. 4q). Neutrophils tended to colonize more T cell^rich^ areas in macrophage-depleted (CD64^DTR^) tumors when compared to controls or macrophage-depleted tumors treated with ICG-001, even though the variability in neutrophil infiltration compromised statistical significance. Importantly, the reduced tumor coverage by neutrophils compared to T cells was not due to insufficient numbers of neutrophils that infiltrated the tumor as T cell and neutrophil numbers were comparable (Fig. S2a, 4i). This demonstrates that some fibrillar regions present topographies permissive to both T cells and neutrophils, while others are specifically permissive to T cells, consistent with our “neutrophil-trained” random forest model not being able to predict T^rich^ regions (Fig. 4r).

Overall, our work shows that fibrillar collagen topographical features are sufficient to predict the localization of T cells and neutrophil infiltration in the tumors. However, while the recruitment of both cell types is dependent on collagen fibers and collagen 3, T cells and neutrophils do not share all topographical determinants, leading to colonization of different regions harboring specific topographies.

## Discussion

We here show that macrophage depletion at late stages of tumorigenesis results in a drastic remodeling of fibrillar collagen in colorectal adenocarcinoma subcutaneous tumors, associated to an increased permissiveness of the tumor to immune infiltration. Specifically, we demonstrated that macrophage depletion leads to a switch from an aligned network of fibrillar collagen, reminiscent of TACS-2, to a network of intermingled fibers with no dominant orientation, resembling TACS-6. This rearrangement relies on an increased deposition of collagen 3, under the control of the transcription factor Tcf4, in both fibroblasts and cancer cells. Most importantly, topographical features of the collagen networks alone are sufficient to predict local accumulation of both T cells and neutrophils.

Our work establishes ECM fine topography as a major determinant of immune landscape in tumors at two levels: the penetrance of immune cells in the tumor core, and their subsequent local accumulation in specific areas. A dense, crosslinked collagen capsula at the tumor edge has long been considered a key barrier restricting T cell infiltration, with T cells accumulating at its periphery^12,16,53,54^. Previous studies demonstrated that disrupting the capsula integrity by degrading collagen *ex vivo* or inhibiting its crosslinking *in vivo* enables T cells to penetrate deeper into the tumor core^12,53^. Our results expand upon this concept, showing that macrophage depletion alters the collagen topography, effectively reshaping the tumor extracellular matrix into a non-canonical fibrillar collagen network enriched in collagen 3. This reorientation of collagen fibers appears to create new migratory paths for T cells to infiltrate the tumor, as supported by our *in vitro* experiments on synthetic fibers. While macrophages have been shown to retain T cells at the tumor edge by directly interacting with them^55^, our results hence indicate that they also play an indirect role by shaping ECM architecture. Interestingly, this permissive topography not only facilitates T cell migration but also enables neutrophil infiltration, revealing a broader immune accessibility.

Machine learning analysis further revealed that the specific accumulation of T cells and neutrophils within collagen-rich regions can be predicted by fibrillar collagen topographical features alone. While some favorable topographies are shared between T cells and neutrophils, other collagen architectures are specifically associated with T cells, suggesting a broader spectrum of favorable networks for T cells compared to neutrophils. Furthermore, the predictive topographical features for T cells differ between control and macrophage-depleted tumors. This points to either the emergence of new collagen topographies permissive to T cells or to a shift in their accessibility due to additional local cues such as chemokines. Deciphering the precise contribution of ECM topography versus chemokine-driven recruitment remains a challenge, as many chemokines are ECM-bound^56–58^. The emergence of reductionist experimental approaches integrating multiple guidance cues should help address this central question.

Through the causative algorithmic (MIIC) analysis of the tumor transcriptome, we shed light on a pro-fibrotic pathway likely resulting in collagen chaotic organization. Macrophage depletion leads to the upregulation of Tcf4 expression in cancer cells and fibroblasts, leading to an increased collagen 3 deposition, which has been reported to promote fibrillar collagen network disorganization^39,40^. Inhibiting Tcf4 in macrophage-depleted tumors rescued the control phenotype. Our results differ from other studies on the MC38 orthotopic model and on spontaneous breast tumors, where macrophage removal rather led to an impaired fibrillogenesis through a decrease in collagens and crosslinking enzymes production by fibroblasts and macrophages themselves. These effects were all mediated by the deprival of macrophage-derived TGF-β. While TGF-β is widely recognized as a master regulator of fibrosis, our findings shed light on an alternative pro-fibrotic pathway mediated by Tcf4. Tcf4 has been previously reported as profibrotic in the liver^51^, but to the best of our knowledge its role in cancer-associated fibrosis has not been reported yet. Importantly, we do not exclude a contribution of TGF-β in our system, potentially through a direct antagonism with the Tcf4 pathway, as reported *in vitro* in fibroblasts^59^. Overall, our findings expand the understanding of fibrosis mechanisms governed by macrophages in the tumor microenvironment, highlighting the critical role of *Tcf4* and challenging the traditional dominance of TGF-β as the main driver of fibrosis in tumors.

Noteworthy, we observed the same transcriptional pro-fibrotic axis in both fibroblasts and cancer cells, suggesting that the Tcf4-Col3a1 pathway that we highlight could be a common trait of mesenchymal cells such as fibroblasts and the here-used MC38 tumor model, consistent with Tcf4 description as a pro-epithelial-to-mesenchymal transition transcription factor^60^. This is consistent with the C2 vs C1 signature correlating with Tcf4 and Col3 in tumoral but also in juxta tumoral tissues, which suggests that the Tcf4-Col3a1 fibrotic axis is at play in non-tumoral tissues. EMT-related transcription factors such as Tcf4 can therefore act as double-edged swords. On the one side, they may promote immune infiltration as reported here, but on the other they support cancer cell survival and invasion. For example, in immunodeficient mice, Tcf4 inhibition leads to a reduction in tumor growth due to increased apoptosis of cancer cell^61,62^. The shift in ECM composition and increase in collagen 3 content could also modulate the balance between tumor growth and T cell activity. Collagen 3 has been shown to be cytostatic in some cancer models^39,40,63^ while the interaction of T cells with collagen through inhibitory receptor LAIR-1 was reported to favor T cell exhaustion^64^. The exhausted profile of CD8^+^ T cells and their interactions with other immunomodulatory immune populations including neutrophils can also render them inefficient at tumor cell killing^65,66^. Surprisingly, macrophage depletion did not impact T cell transcriptional profiles and T cell subset proportions which could result in unchanged dynamics between inhibitory and stimulatory signals/T cells. The pro- and anti-tumoral consequences of macrophage depletion could therefore balance each other, preventing tumor reduction despite a massive infiltration of T cells as we describe here. This could explain the poor clinical benefits of therapeutic strategies targeting macrophages and EMT pathways, including CSF1R and β-catenin/Tcf4 inhibition^67–69^. According to our results, failure of such therapies could result from an increase in EMT pathways promoting cancer cell growth in anti-CSF1R therapies, and a decrease of T cell infiltration/responses in therapies targeting β-catenin/Tcf4 complex.

In conclusion, our findings uncover a novel interplay between macrophages, EMT pathways and tumor ECM topography. We established that macrophage depletion induces a transition from a canonical aligned collagen TACS-2 capsula to an intermingled collagen-3-rich TACS-6-like network, promoting immune infiltration of the tumor. Our work supports recent publications aiming at using collagens and TACS as prognosis markers^70–72^. Yet, this remains challenging, as diverse TACS tend to have dual effects on tumor and immune cells: TACS-2 would exclude T cells from the tumor but prevent tumor cells from exiting the tumor mass and form metastasis^13,14^, TACS-6 would promote immune infiltration but also allow multidirectional tumor cell migration^13^. Unraveling how the diverse TACS are established and evolve over the duration of tumorigenesis and characterizing their effects on both cancer and immune cells is essential to better understand the tug-of-war between the tumor and the immune system. This fundamental knowledge will open new avenues to improve immunotherapies and anti-EMT drugs but also design new therapies directly targeting the ECM.

## Materials and Methods

### Mice

All mice experiments were performed in accordance to the European and French National Regulation for the Protection of Vertebrate Animals used for Experimental and other Scientific Purposes (Directive 2010/63; French Decree 2013-118, Authorization DAP number APAFIS #36924-2022042210525004 v2 given by National Authority). Mice were bred and maintained under specific pathogen-free conditions in our animal facilities of Institut Curie, following institutional guidelines, and housed in a 12-h light/12-h dark environment with free access to water (osmotic water) and food (MUCEDOLA, 4RF25SV Aliment Pellets 8 × 16 mm irradiated 2.5 Mrad. CD64-hDTR mice^34^ were provided by Sandrine Henri (CIPHE, Marseille, France) and crossed on C57BL6/J mice obtained from The Jackson Laboratories. tdTomato/GFP^73^ mice were provided by Lequn Luo (Stanford, USA). Experiments were performed on 8- to 14-week-old male or female mice in our animal facility under the approval number D750617. Each *in vivo* experiment was conducted on at least 4 mice per experimental group combining males and females and using littermates or sex- and age-matched mice as controls without randomization or blinding. Mice showing physical signs of inflammation, infections or unethical weight loss were excluded from experiments.

### Cells

MC38 cells were provided by Nicole Haynes (Peter MacCallum Cancer Centre, Melbourne, Australia). MC38 cells were cultivated in RPMI Glutamax (Gibco, 61870036), supplemented with 10% of Fetal Bovine Serum (Biowest, S1810), 1% Penicilline/Streptomycine (ThermoFisher, 15070063), and 0.1% b-mercaptoethanol (Gibco, 11528926). Cells are seeded at 0.2x10^6^ cells / mL and passaged 1:4 every 2-3 days when reaching confluence. Cells are adherent and thus detached from culture plates using TrypLE™ Express (Gibco, 12605010). Cells were always passaged to 1:3 one day prior to injection in the mice to reach 60% confluence before detachment for *in vivo* injection.

### In vivo tumor growth and macrophage depletion

8 to 14-week-old CD64 WT (CD64^WT^) and CD64-hDTR (CD64^DTR^) mice (males and females) were injected with 0,5x10^6^ MC38 cells in 100 µl in Phosphate Buffer Saline (PBS) subcutaneously. They were then injected intraperitoneally (i.p.) with 50 µg/kg of diphteria toxin (Merck,322326-1MG) in PBS at day 9 d.p.i. and then 17,5 µg/kg at day 11 d.p.i. and 13 d.p.i., and sacrificed at day 14 d.p.i. Tumor growth was analyzed using a digital caliper and tumor volume was calculated using the following formula: (length*width^2^)/2.

### Tcf4 in vivo inhibition

CD64^WT^ and CD64^DTR^ mice were injected i.p. with 10 mg/kg of ICG-001 (MedChem, HY-14428), resuspended in DMSO (Fisher, D139-1) and diluted in 30% Polyethylene glycol (Sigma, 81169) 30% 1-2, Propanediol (Sigma, 16033)) at day 9, 11 and 13 d.p.i. concomitantly with DT injections, and sacrificed at day 14 d.p.i.. Control mice were injected with the same amount of DMSO diluted in 30% Polyethylene glycol, 30% 1-2, Propanediol.

### Primary T cell isolation and activation

Splenocytes of tdTomato/GFP male and female mice were harvested and T cells were isolated using dynabeads untouched Pan t cell isolation kit (Dynabeads,10425073) following manufacturer’s instructions. Isolated T cells were then activated using CD3/CD28 beads (Gibco, 10022593, 2 beads for 1 T cell) and 25U/ml of IL-2 (Novartis, Proleukin) in T cell media (RPMI (Gibco, 61870-010), 10% FBS (Biowest, S1810-500), 1% Penicillin/Streptomycin (Eurobio, CABPES01-0U), 1% HEPES(Gibco, 15630-056), and 1% non-essential amino acids (Gibco,11140-035). After 3 to 5 days, beads were washed off the cell using a magnet and rinced twice with T cell media.

### In vitro seeding of primary T cells on fabricated 3D fibers

Deformable fiber arrays were fabricated using TPP with a QSwitch Teem Photonics laser (10 kHz, 5 ns pulses, 532 nm, 10 μJ) and a IX70 Olympus microscope equipped with water (60× NA 1.2) and oil (100× NA 1.4) objectives, a piezo-z stage, and a Guppy CCD camera. The system was controlled by Lithos software with autofocus functionality.

Fibrous scaffolds showing aligned or intermingled topographies were designed using Sympoly V4. Each scaffold consisted of three 21-fiber layers (160 μm length, 8 μm z-spacing). Slicing parameters included a lateral voxel size of 0.17 μm and a vertical voxel size of 0.55 μm for fibers (overlap: 80%). Two resins were used: Resin 1 (PEGDA575 (Sigma, 26750-48-9)) with 15% PETA (Sigma, 986-89-4) and 5% Irgacure 651 (Sigma, 24650-42-8) for anti-adhesive supporting walls, and Resin 2 (PEGDA250 (Sigma, 25852-47-5) with 10% PETA and 5% Irgacure 651) for fibers. Supporting walls were printed with a 60x objective (laser power: 6.4 mW, exposure: 1000 μs) and Resin 2 was added to print fibers using a 100× objective (laser power: 4.9 mW, exposure: 800 μs). Scaffolds were washed in ethanol and coated with 10 μg/mL fibronectin (Thermofisher, 33010018) labeled with CF™640R (Sigma, SCJ4600044). Scaffolds were washed in RPMI medium before seeding 100,000 activated T cells (0.5 mL) at a density of 15–30 cells per scaffold. Cells were allowed to settle for 20 minutes at 37°C and 5% CO₂. Live imaging was performed on two spinning-disk confocal microscopes (Nikon Eclipse Ti-E, CSU-X1, Yokogawa). Images were acquired with a 561 nm laser for T cells and a 633 nm laser for fibers using Prime BSI or QuantEM cameras. Image stacks (180×280×31 μm, voxel size: 0.16×0.16×1 μm or 0.4×0.4×1 μm) were collected over 5 hours at ΔT = 2.5–3 min intervals. For the analysis, Z-planes were selected in ImageJ (version from May 2017, provided by NIH, Maryland, USA) to exclude cells on glass, and a median filter (1 μm) was applied. T cell tracking was performed using IMARIS (v9.2) with spot detection (size: 10–12 μm) and a linking max distance of 20–30 μm. Data were processed in MATLAB (R2022b) to compute 2D velocity and distance ratios over a 1-hour period.

### Immunoffuorescence

Tumors were collected in 4% paraformaldehyde (Edimex, 3178-200-19), incubated for 24h at 4°C in obscurity. They were dehydrated in two successive baths of 15% and 30% sucrose (Fisher Scientific, 10634932), included in Optimum Cutting Temperature (TissueTek, 4583) and stored at -20°C. Tumors were cut into 20 µm sections for SHG imaging, or 7 µm sections for epifluorescence imaging. Sections were permeabilized using 0,2% Triton (Sigma, T8787) diluted PBS, blocked using 1:100 Fcblock (BD, 553142, 1:100), 1:20 Goat serum (JacksonImmunoResearch, 005-000-121) or Donkey serum (JacksonImmunoResearch, 0017-000-121) depending on the host of secondary antibodies, and 3% IgG-free Bovine Serum Albumin (JacksonImmunoResearch,001-000-162) in 0,05% Triton in PBS. They were stained with the following primary antibodies: anti-F4/80-AF647 (Biorad, MCA497A647, 1:100), anti-CD3e (Biolegend, 100306, 1:200), anti-Ly6G (Biolegend, 127605, 1:200), anti-Col3a1 (SouthernBiotech,1330-01, 1:300), anti-Tcf4 (ProteinTech, 16801875, 1:100). For Col3a1 and Tcf4 stainings, donkey anti-goat AF647 (Invitrogen, A21202, 1:200) and donkey anti-rabbit AF546 (Invitrogen, A10040, 1:00) secondary antibodies were used respectively. Slides were mounted in Fluoromount-G™ with or without DAPI (Invitrogen™, 15596276 or 15586276 resp.).

### Image acquisition by epiffuorescence and two-photon microscopy

Epifluorescence imaging was performed using an AxioScan Z1 microscope (Zeiss) at 20X (Zeiss Plan-Apochromat 20x/0.8 #420650-9901-000). Two-photon imaging was performed using an inverted two-photon laser-scanning confocal microscope (Leica SP8), coupled with a femtosecond laser (Chameleon Vision II, Coherent Inc.) using a 20×/0,75W HC IRAPO water immersion objective. The excitation wavelength was set at 940 nm and signals were acquired using 3 non-descanned HyD detectors: 525/40 nm (GFP), 585/40 nm (tdTomato) and <492 nm (SHG). The acquisition was performed in resonant mode, with Z-size of 10 µm and a Z-step of 2 µm, and tiling arrays were performed on the whole tumor. Image stitching was performed using the LAS X software (Leica).

### Picrosirius red staining

2D sections of the tumors (5 µm) were collected and stained in Picro Sirius Red solution for 60 minutes (Sigma, 365548) to stain for collagen content. Coverslides were then washed, mounted, and analyzed after scanning with a Nanozoomer Hamamatsu (MicroPiCell Facility, Nantes University, France).

### 3D reconstruction of tumors by Kratoscope autoffuorescence imaging

Tumors were harvested, fixed in 4% paraformaldehyde overnight at 4°C and subsequently dehydrated in successive increasing concentration of ethanol baths. Samples were then embedded in 5% Oil Red-O-stained paraffin (Diapath, C0512) as described in Malloci *et al.*^73^. Serial block face imaging of the samples was performed using the Kratoscope system (Kaer Labs, France) to detect native tissue autofluorescence. Briefly, the tumors were mounted on an automated microtome and sectioned into 10 µm thick sections. The tissue block was imaged using a 405nm excitation LED. The successive imaging-sectioning cycles generated a stack of images reflecting the autofluorescence throughout the whole sample and was used to perform 3D reconstruction of the total tumor volume. Individual 2D sections were collected during the process for downstream histological staining.

### Orion multiplex imaging

#### • Antibody Preparation and Conjugation

All antibodies were monoclonal, rabbit-derived, and carrier-free. The MHCII antibody was directly conjugated to biotin, while other antibodies were conjugated to ArgoFluor dyes using amine conjugation chemistry (Table 1).

**Table 1:**
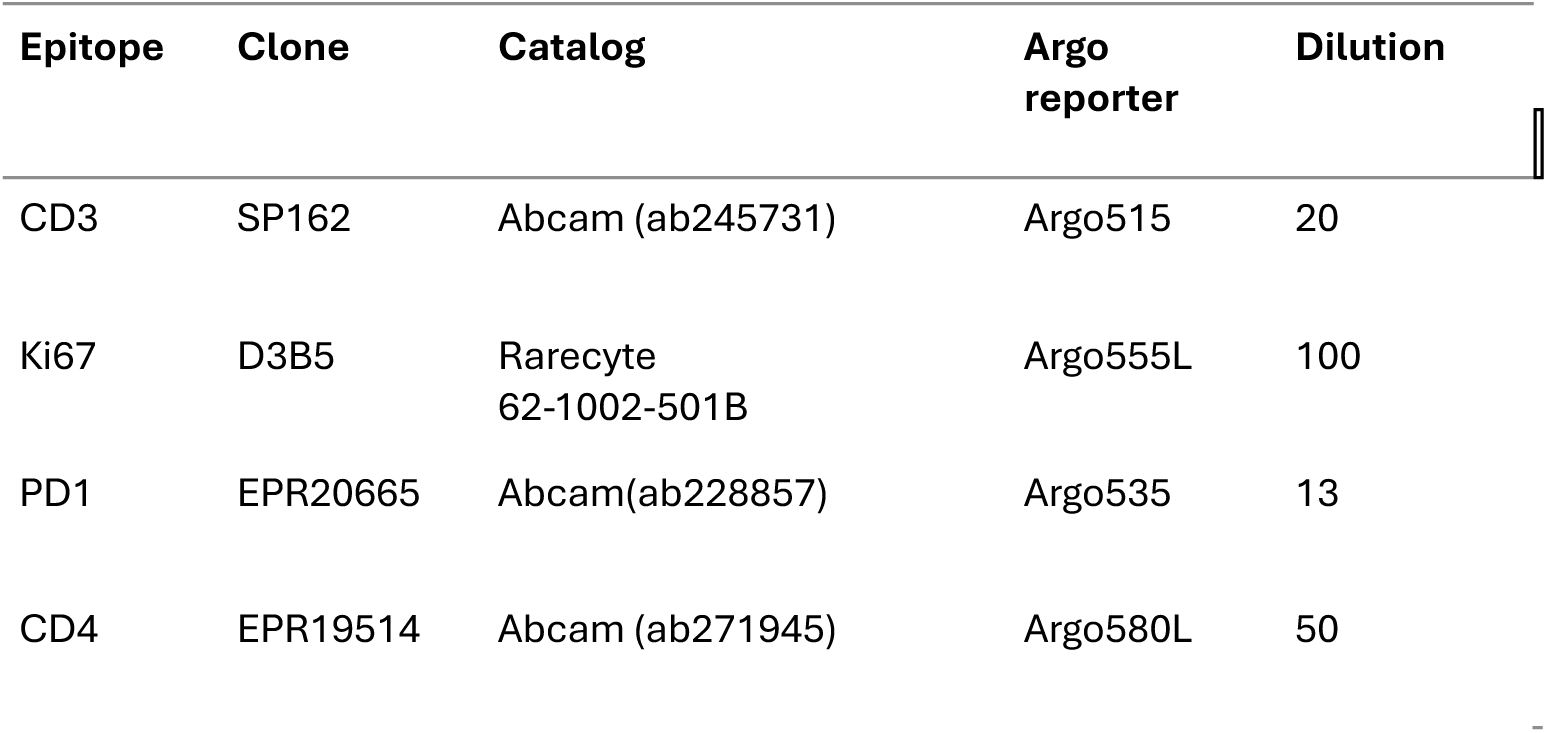
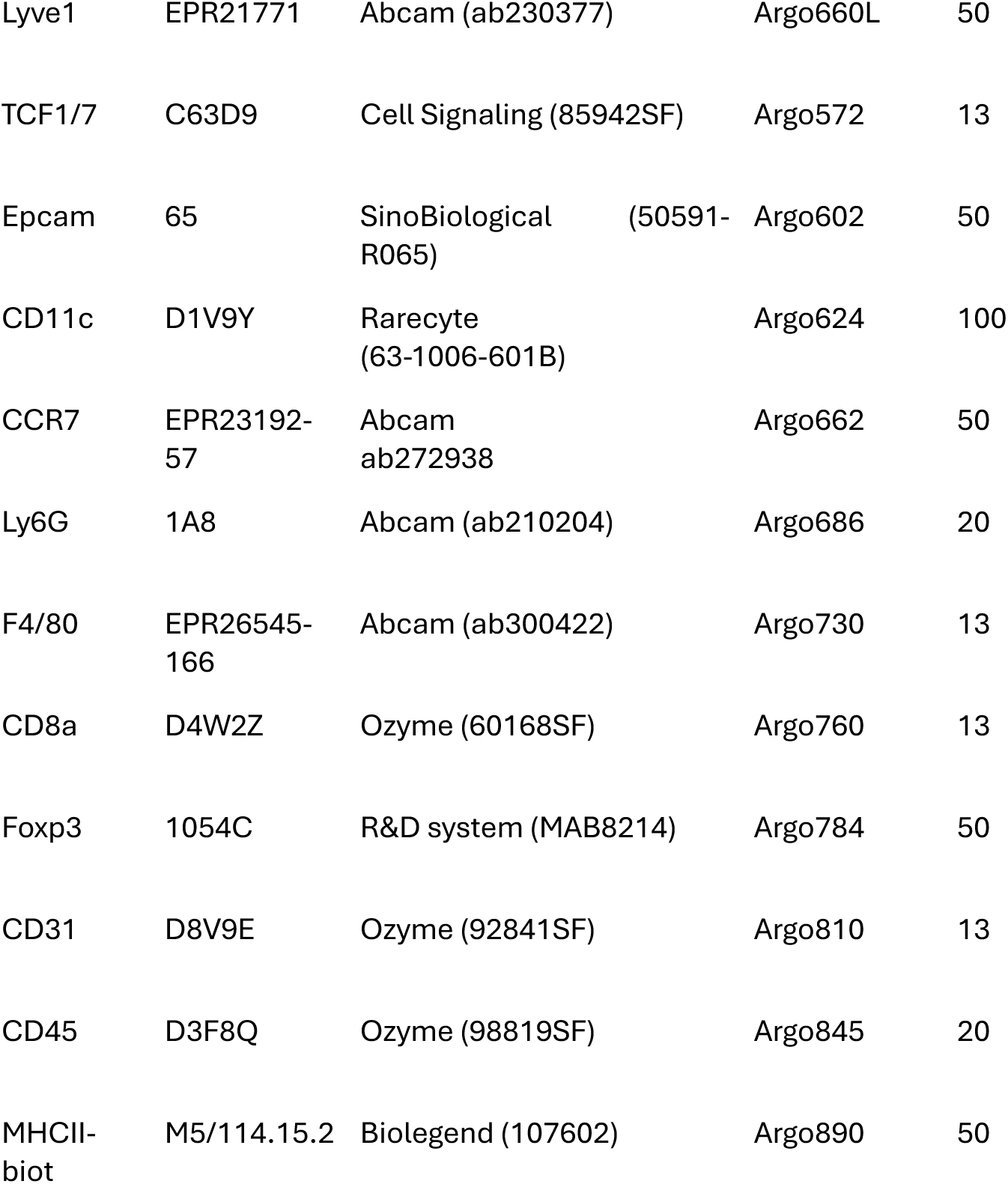
Primary antibodies panel conjugated with ArgoFluor.

| Epitope | Clone | Catalog | Argo reporter | Dilution |
| --- | --- | --- | --- | --- |
| CD3 | SP162 | Abcam (ab245731) | Argo515 | 20 |
| Ki67 | D3B5 | Rarecyte<br>62-1002-501B | Argo555L | 100 |
| PD1 | EPR20665 | Abcam(ab228857) | Argo535 | 13 |
| CD4 | EPR19514 | Abcam (ab271945) | Argo580L | 50 |
| Lyve1 | EPR21771 | Abcam (ab230377) | Argo660L | 50 |
| TCF1/7 | C63D9 | Cell Signaling (85942SF) | Argo572 | 13 |
| Epcam | 65 | SinoBiological (50591-R065) | Argo602 | 50 |
| CD11c | D1V9Y | Rarecyte (63-1006-601B) | Argo624 | 100 |
| CCR7 | EPR23192-57 | Abcam ab272938 | Argo662 | 50 |
| Ly6G | 1A8 | Abcam (ab210204) | Argo686 | 20 |
| F4/80 | EPR26545-166 | Abcam (ab300422) | Argo730 | 13 |
| CD8a | D4W2Z | Ozyme (60168SF) | Argo760 | 13 |
| Foxp3 | 1054C | R&D system (MAB8214) | Argo784 | 50 |
| CD31 | D8V9E | Ozyme (92841SF) | Argo810 | 13 |
| CD45 | D3F8Q | Ozyme (98819SF) | Argo845 | 20 |
| MHCII-biot | M5/114.15.2 | Biolegend (107602) | Argo890 | 50 |

Antibodies were first concentrated by filtering on Amicon filters (Sigma-Aldrich, UFC503096). Following centrifugation at 14,000 × g for 10 minutes, 475 µL of Reaction Buffer (Sigma-Aldrich, 36486) was added. The solution was transferred to a clean Amicon collection tube and centrifuged at 1,000 × g for 2 minutes, yielding 25 µL of concentrated antibody (2-5 mg/mL). To conjugate the antibodies, 5 µL of ArgoFluor dye dissolved in DMSO was added to the concentrated antibody solution and vortexed for 10 seconds and incubated in the dark at room temperature for 1 hour. The reaction was quenched by adding 1.5 µL of 1 M Tris-HCl (Teknova, T1083), followed by vortexing on high for 5 seconds and further incubation in the dark for 20 minutes at room temperature. The conjugate was purified using a clean Zeba column (ThermoFisher, 7766) by centrifugation at 5,000 × g for 5 minutes. The final volume was measured, and labeling efficiency was determined via absorbance spectroscopy. The conjugated antibodies were diluted in PBS Antibody Stabilizer (Candor Bioscience, 131125) to a concentration of 0.1 mg/mL.

#### • Slide Preparation

Slides were immersed in a bleaching buffer (30% H2O2, 1 M NaOH in PBS) and exposed to LED light for 1 hour to quench autofluorescence. Subsequently, slides were transferred to a UV transilluminator chamber (365 nm, high setting) for 30 minutes. Excess bleaching buffer was removed, and slides were washed twice with PBS (5 minutes each).

Slides were washed twice with PBS-T (25% Triton) and twice with PBS (5 minutes each). In a humidity chamber, slides were incubated with Image-iT FX Signal Enhancer (Invitrogen, I36933) for 15 minutes, followed by two washes with PBS-T (25% Triton) and one wash with PBS (5 minutes each). A cocktail solution of all conjugated antibodies (Table 1) was prepared in PBS Antibody Stabilizer (Candor Bioscience, 131125) with 10% rabbit serum. Slides were incubated with the antibody cocktail in a humidity chamber for 1 hour and 30 minutes in the dark at room temperature. Slides were then washed twice with PBS-T (25% Triton) and once with PBS (5 minutes each). A secondary staining solution was prepared with PBS Antibody Stabilizer (Candor Bioscience, 131125), 10% goat serum, 0.1% Hoechst 33342 (Thermo Fisher Scientific, H3570), and 1% Streptavidin ArgoFluor 890 (Rarecyte, 52-1026-802). Slides were incubated with the secondary solution in a humidity chamber for 30 minutes in the dark at room temperature. Slides were then washed once with PBS-T (25% Triton) and once with PBS (5 minutes each). Slides were coverslipped with ArgoFluor Mounting Medium (Rarecyte, 42-1214-000) and left to dry overnight.

#### • Image acquisition

Images were acquired at 20× magnification using the Orion imaging system (Rarecyte) and processed with Artemis software (version 4.1.0.28). Assembled OME.TIFF files were generated for each slide.

### Image analysis

Regional analysis were performed by dividing whole tumor images into 100 µm x 100 µm tiles and extracting cell number or staining densities or intensities using QuPath software (v0.4.4). Subsequent analysis was performed using R software (v4.3.1). For each tile, cell or staining enrichment was determined as the cell or staining density (threshold determined using QuPath) was superior to the mean cell or staining density of positive regions (cell number or staining density > 0).

Solidity measurements was performed using Fiij software. Briefly, tumor masks were drawn by hand (delineating the whole tissue and removing the skin), and the shape descriptor plug-in was applied.

### GLCM Analysis

The gray level co-occurrence matrix^74^(GLCM) algorithm was applied to characterize the texture of collagen. For each sample, a minimum of 5 regions (318 µm × 318 µm) from the core and border were manually selected. For each region, the GLCM was computed, and textural homogeneity was extracted. The gray level co-occurrence matrices and homogeneity properties were calculated using functions from the Scikit-Image Python library^75^: skimage.feature.graycomatrix and skimage.feature.graycoprops, respectively. The GLCM, with 256 intensity levels, was computed with the following parameters: distances from 1 to 100 pixels with a step of 1 pixel (1 pixel = 0.45µm), and angles of 0°, 45°, 90°, and 135°.

### Topographical feature extraction from SHG images

SHG images were analyzed using the OrientationJ plug-in (developed by EPFL, Switzerland) within the ImageJ software . This function computes the structure tensor for every Gaussian-shaped window (with a size of 20-60-100-140 pixels) by determining the continuous spatial derivatives along the x and y axes through Gaussian interpolation. The structure tensor is a symmetric matrix computed in regions spaced 40 pixels one from the other. The tensor can be characterized by three quantities (derived from the two eigenvectors and eigenvalues): the energy (trace of the tensor, related to the size of the ellipsis associated to the tensor), the coherency (difference of the eigenvalues normalized by the trace, related to the aspect ratio of the ellipsis) and the orientation (angle formed by the major axis of the ellipsis with the x-axis of the image). Intensity mean and variance, density and density variance, and distance of the region from the border were extracted using ImageJ software. The number of cells (either Ly6G^+^ or CD3ɛ^+^) in 100 µm x 100 µm windows were extracted from Qupath software ^76^. Principal Component Analysis was performed using *prcomp* function in R (v4.3.1) software.

### Random forest classification

Random Forest predictions were performed using a custom-made Python code. 70% of the initial data was used for training the model, and the remaining 30% for the test set. A stratified 10-fold cross-validation was performed on the training set, during which a RandomSearch (N=50) was conducted to find the best combination of parameters based on the performance obtained on the validation set.

The complete R-code is available *here*.

For evaluating random forest performances, we use various metrics based on the true positives (TP), true negatives (TN), false positive (FP) and false negative (FN) which are derived from the predicted versus observed target class.

Accuracy is the proportion of correctly classified instances out of the total instances, formulated as: Accuracy = (TP + TN) / (TP + TN + FP + FN).

Balanced accuracy corresponds to the macro-average of the recall score per class, here it is formulated as: Balanced Accuracy = (1/2) x (TP / (TP + FN) + TN / (TN + FP)),

The Precision corresponds to the proportion of true positive out of all positive predicted instances denoted by: Precision = TP / (TP + FP),

The Recall measures the proportion of true positive out of all observed positive instances where: Recall = TP / (TP + FN),

The Area Under the Receiver Operating Characteristic Curve score (ROC AUC) denoted as AUC is a measure of the model’s ability to discriminate between positive and negative classes across different probability thresholds.

The Matthews correlation coefficient (MCC) takes into account the TP, TN, FP and FN rate and is especially useful for imbalanced datasets. It is formulated as:

MCC = (TN x TP - FN x FP) / square-root ((TP + FP) x (TP + FN) x (TN + FP) x (TN + FN)).

The F1 score corresponds to the harmonic mean of precision and recall, providing a balance between the two metrics, with: F1 = (2 × Precision × Recall) / (Precision + Recall).

### FACS analysis of the blood

80 µL of blood was collected submandibularly in 10 µL of 100mM EDTA (ThermoFisher Scientific, 15575020) diluted in PBS and centrifuged 10 min at 2000xg at 4°C. The pellet was resuspended in 7 mL of RBC lysis buffer 1X diluted in water and incubated on ice for 5 min. After adding at least 7 mL of FACS buffer, centrifugation 7 min at 1500rpm, cells were resuspended in FACS buffer and plated in a 96 well plate. After centrifugation (5min, 1500rpm), cells were incubated 10 min with Fcblock solution diluted in FACS buffer (BD, 553142, 1:100), washed with FACS buffer and incubated with following antibodies : anti-SiglecF PE/Dazzle 594 (Biolegend, 155530,1:400), anti-CD115 APC-Cy7 (Biolegend, 135531, 1:200), anti-GR1 BV510 (BD, 563040, 1:200), anti-B220 PE (BD, 553090, 1:200), anti-CD4 FITC (BD, 553047, 1:300), anti-CD8 APC (BD, 553035, 1:300), anti-NK1.1 PerCP-Cy 5.5 (BD, 551114, 1:200). After washing with FACS buffer, 10 µl of counting beads (Invitrogen, 01-1111-42) were added. Flow cytometry was performed on an Aurora Analyzer (Cytek) and analyzed with FlowJo software (version 10.8.1).

### Atomic force microscopy

30 μm thick cryosections were prepared at -20°C from tumors embedded in OCT compound and immediately transferred to a poly-L-lysine coated glass-bottom dish. PBS containing Complete (TM) EDTA-free Protease Inhibitor (Sigma) and 10% horse serum was added on top of the sections as soon as they melted to avoid desiccation. CD3e staining was then performed in PBS with protease inhibitors, 10% horse serum and 0.2% triton X-100 for one hour at room temperature. The atomic force microscopy measurements were done on the same day.

Atomic force microscopy analysis was performed on A JPK NanoWizard II system (Bruker Nano GmbH) with a CellHesion module mounted on a Zeiss LSM510 confocal microscope (Carl Zeiss NTS Ltd.). We used silicon nitride cantilevers (spring constant: 0.06 N/m, spherical 4.5µm diameter tip; Novascan Technologies). The cantilever spring constant and deflection sensitivity were calibrated in fluid via the thermal noise method (Hutter and Bechhoefer, 1993). We used a 10x objective connected to a webcam attached to the microscope to visualize the samples and to determine the regions of interest. In each location, indentations distributed in a 5×5 points grid (25 µm×25 µm) were performed. The elastic modulus for each force curve was calculated using JPK software (JPK DP version 4.2) assuming a Hertz model of impact.

### Single cell RNA sequencing

MC38 tumors from CD64^DTR^ and CD64^WT^ were harvested at day 14 d.p.i., digested in Liberase TL 0,25mg/ml (Roche #5401020001,) DNAse 0,5mg/ml (Sigma, DN25-100MG) in CO_2_ independent medium (Gibco, 18045088) using GentleMacs Octo dissociator pre-set program (37C_m_TDK1, 41min at 37° under agitation). Cells were filtered on a 100 µm cell strainer and red blood cells were lysed using red blood cells lysis buffer 1X (Biolegend, 420301) diluted in water. After centrifugation (5min, 1500rpm), cells were incubated 10 min with Fcblock solution diluted in FACS buffer (BD, 553142, 1:100) and stained either with 1) DAPI (Invitrogen, 11534886, 1:10000) for the first experiment (batch #1), or 2) with DAPI, anti-CD45 APC-Cy7 (BD, 557659, 1:200) and anti-CD64 APC (Biolegend,139306,

1:200) (for the second experiment (batch #2) diluted in FACS buffer (PBS 1X, 0,5%BSA, 2mM EDTA). For batch#1, DAPI-cells were sorted while for batch#2, DAPI^-^CD45^-^, DAPI^-^ CD45^+^CD64^+^, DAPI^-^ CD45^+^CD64^-^ populations were sorted separately. For batch #1, 15 000 cells form CD64^WT^ and CD64^DTR^ tumors were loaded. For Batch #2, 10000 cells form CD64^WT^ and CD64^DTR^ tumors were loaded. ScRNAseq data were analyzed with R using the Seurat suite (v4). Cells were then filtered out when expressing less than 1000 genes, or when expressing more than 10% of mitochondrial genes. Doublets were identified using the DoubletFinder package^77^. Clusters containing less than 100 cells were excluded from the analysis. Differential analysis was done using the DESeq2 package^52^ and gene set enrichment analysis was performed using the FGSEA package^78^. For sub-clustering of non-immune, Mo/Mac/DC and Lymphocyte clusters, batch #1 and batch #2 CD45^-^, CD45^+^CD64^+^ and CD45^+^CD64^-^, or CD45^+^CD64^-^ sorted samples were selected respectively. Data management, quality control and primary analysis were performed by the Bioinformatics platform of the Institut Curie. Raw counts are available *<u>here</u>*.

### MIIC Analysis

MIIC network inference was performed on non-immune cells. Further processing of the scRNAseq data led to a total of 16737 non-immune cells (Fig. 3a), corresponding to 3303 fibroblasts and 13434 cancer cells including 8838 cancer cells in C1 (Lgals3) cluster and 4566 cancer cells in C2 (Airn) cluster. For the analysis of Cancer cells (C1+C2), a down-sampling to 4000 cells was performed to allow computational calculations.

First, the top 50 and 250 transcription factors with the highest mutual information with the tested condition (CD64^DTR^ vs CD64^WT^) were selected along with the list of collagen genes (44 genes). For cancer cells the number of cells was subsampled to 4000 cells. Full networks are available below:

- For Fibroblasts (CD64^DTR^ vs CD64^WT^) : <u>50 TFs, 250 TFs</u>
- For Cancer cells (CD64^DTR^ vs CD64^WT^) : <u>50 TFs</u>, <u>250 TFs</u>
- For C2 vs C1: 75 TFs + 535 non-TF genes

### Binding motifs enrichment analysis

The 5000 bp upstream of the differentially expressed genes (Log2FC>0.25, adjusted p-value < 0,05) between CD64^DTR^ and CD64^WT^ tumors in fibroblasts and total cancer cells (C1 + C2) or between C2 and C1 independently from macrophage depletion were extracted using the Ensemble browser (https://www.ensembl.org/index.html). Binding motif enrichment analysis was performed on extracted promoter regions using the AME tool from the MEME suite (https://meme-suite.org/meme/tools/ame).

### Human data analysis

Human data from COAD (TGCA, colon adenocarcinoma database) were analyzed using Gepia2 (http://gepia2.cancer-pku.cn/) for Spearman correlation analysis.

### Statistical analysis

Statistical tests used are indicated in the figure legend of each graph. Graphs and statistical analysis were performed using Prism Graphpad (v10.3.1) or R (v4.3.1) softwares. Non-parametrical tests were used due to non-normal distribution of the data. Mann-Whitney test was used to compare two conditions. Two-way-ANOVA was performed when more than two conditions were compared. In all graphs, pooled biological replicates (mice for *in vivo* or cells for *in vitro*) from independent experiments (N) are plotted on the same graphs. N is indicated in each figure legend.

### Materials availability

The authors confirm that the data supporting the findings of this study are available within the article, its supplementary materials or online (links provided within the text). Raw images and flow cytometry data are available upon request to the corresponding authors.

## Acknowledgements

We thank Institut Curie for access to the flow cytometry, animal, and the imaging facilities. The authors greatly acknowledge the Cell and Tissue Imaging (PICT-IBiSA) and the Nikon Imaging Centre @ Institut Curie-CNRS, members of the French National Research Infractucture France-BioImaging (ANR10-INBS-04). Epifluorescence image acquisitions were done at the CHICS core facility (Cordeliers Histology, Imaging, Cytometry and Spatial omics, Centre de Recherche des Cordeliers UMRS 1138, INSERM, Université Paris Cité, Sorbonne Université, Paris, France). We acknowledge the IBISA MicroPICell facility (Biogenouest), member of the national infrastructure France-Bioimaging supported by the French national research agency (ANR-10-INBS-04) for picrosirius red staining and image acquisition. High-throughput sequencing was performed by the ICGex NGS platform of the Institut Curie supported by the grants ANR-10-EQPX-03 (Equipex) and ANR-10-INBS-09-08 (France Génomique Consortium) from the Agence Nationale de la Recherche ("Investissements d’Avenir" program), by the ITMO-Cancer Aviesan (Plan Cancer III) and by the SiRIC-Curie program (SiRIC Grant INCa-DGOS-465 and INCa-DGOS-Inserm_12554). The authors wish to thank Sonia Lameiras, Raphaël Narbey and Zacarias Garcia for technical advice and help, and Philippe Bousso for providing equipment, Alexandre Boissonnas, Emmanuel Donnadieu, Louise Dupuis, Adrianna Lecourieux and the Lennon-Duménil team for discussions and Romane Henninger for comments on the manuscript. ZF was funded by ITMO Cancer. This work was supported by ANR-JCJC *InfEx* (ANR-20-CE15-0023) and ARC PJA1 *NexT* (ARCPJA2023070006800) to HDM, ERC Synergy *SHAPINCELLFATE* to AMLD, the *Fondation Chercher Trouver*, and ANR-10-IDEX-0001-02 PSL (LabEx DCBIOL). This work has received support under the program France 2030 launched by the French Government. FS and HI acknowledge funding from ANR-22-PESN-0002 *AI4scMed*. This work was also supported by the Finnish Cancer Institute (K. Albin Johansson Professorship); Research Council of Finland Centre of Excellence program (# 346131) and ERC AdG *BorderControl* to JI. Work done by GS is supported in part by the following grants: ERC-Synergy (Grant# 101071470); AIRC-IG (Grant#22821); AIRC 5x1000 (#22759); by the Italian Ministry of University and Research (PRIN202223GSCIT_01/G53D23002570006 /20229RM8A_001; COMBINE/G53D23007040001/P2022RH4HH002; PNRR_ CN3RNA _SPOKE/G43C22001320007).

## Authors contributions

Conceptualization: ZF, HDM, AMLD, PP Methodology: ZF, HDM, PP, CG, EM, GS, JCG, JI, HI Investigation: ZF, LC, IC, MM, AC, FPF, EL, VM, LL

Formal analysis: ZF, PP, FS, RJM, MT, JGC Funding acquisition: HDM, AMLD Resources: SC, VS, EP

Visualization: ZF, HDM

Writing – original draft: ZF, HDM

Writing – reviewing and editing: AMLD, ZF, HDM

---

**Supplementary figure 1:**
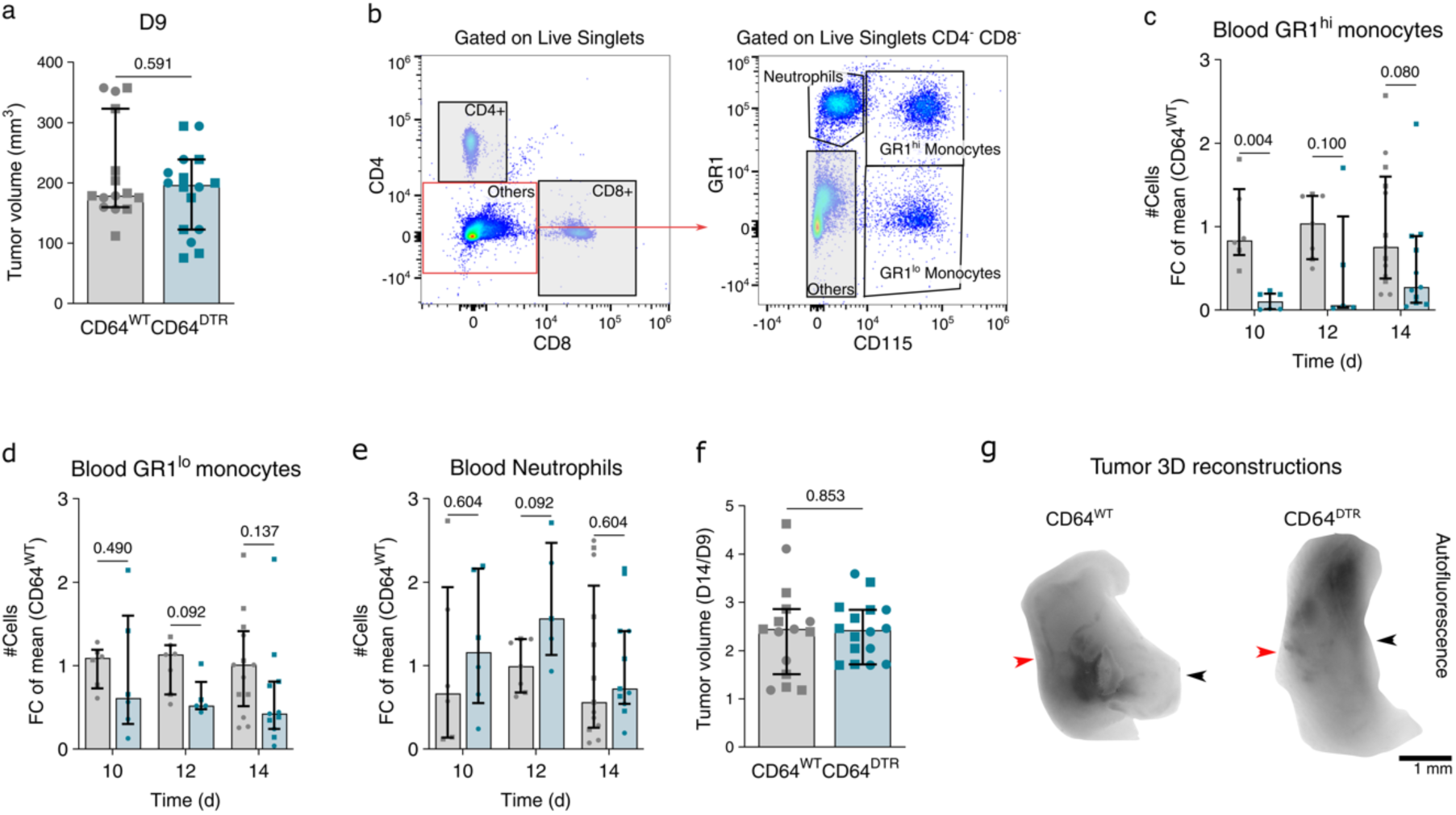
Macrophage depletion disrupts collagen topography at the tumor border. **(a)** Tumor volume of CDC4^WT^ and CDC4^DTR^ tumors at day 9 (N=3 independent experiments; median and interquartile range, Mann-Whitney test). **(b)** Gating strategy of flow cytometry analysis of the blood. **(c)** Number of CD115^high^GR1^high^ monocytes, **(d)** CD115^high^GR1^low^ monocytes, **(e)** and CD115^-^GR1^high^ neutrophils in the blood of CDC4^DTR^ and CDC4^WT^ mice at day 10, 12 and 14 (N=2 independent experiments; median and interquartile range, two-way Anova test). **(f)** Volume of CDC4^WT^ and CDC4^DTR^ tumors at day 14 normalized by their volume at day 9 (N=3 independent experiments; median and interquartile range, Mann-Whitney test). **(g)** Representative autofluorescence images of 3D reconstructions of CDC4^WT^ and CDC4^DTR^ tumors at day 14 (N=1 experiment; black arrow = tumor border; red arrow = skin).

**Supplementary figure 2:**
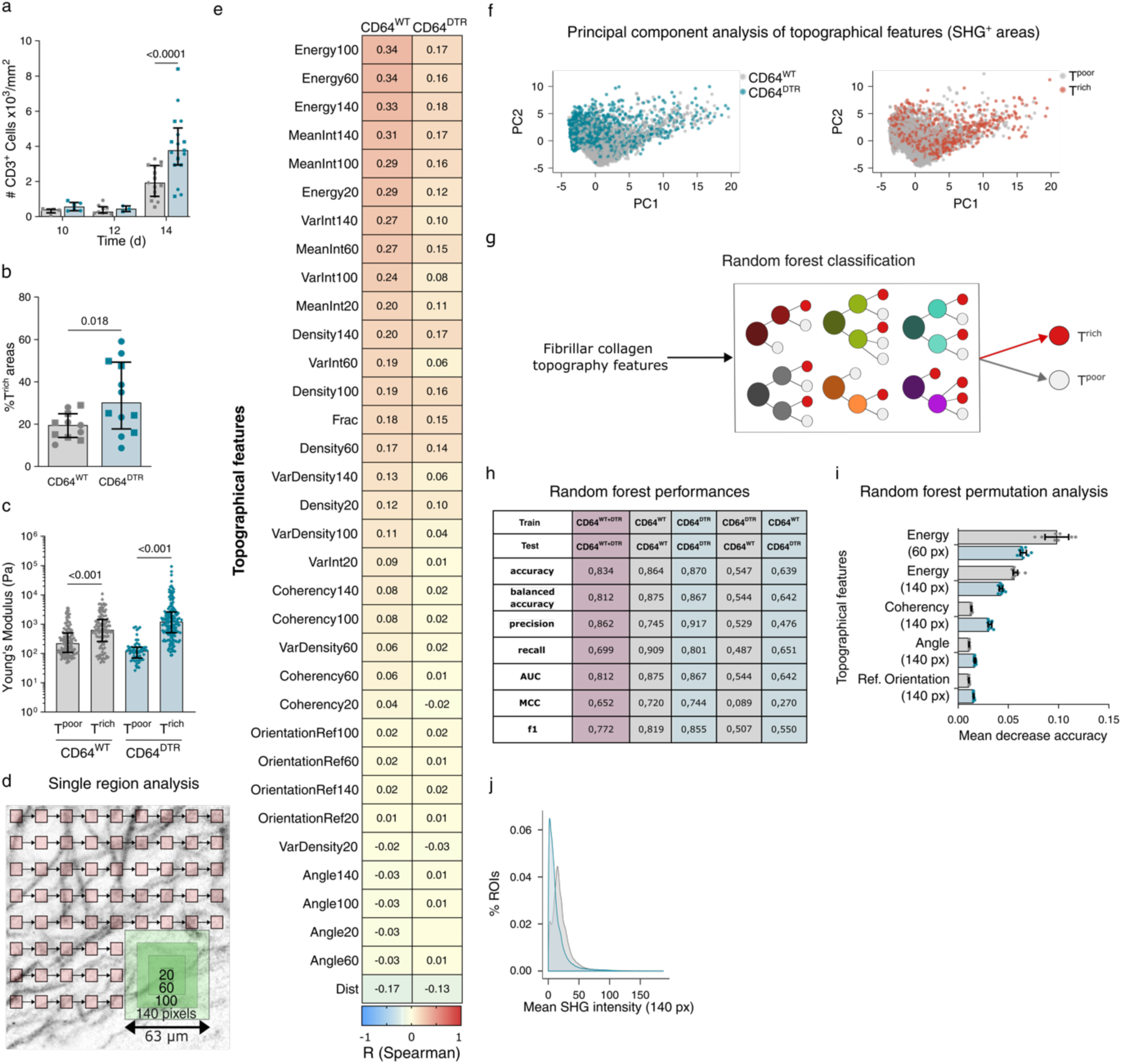
Collagen remodeling is associated to a drastic increase in T cell infiltration in specific network topographies. **(a)** Number of CD3^+^ cells/mm^2^ in CDC4^WT^ and CDC4^DTR^ tumors at day 10,12,14 (N=2 for day 10 and 12; N=3 for day 14; Mann-Whitney test, median and interquartile range). **(b)** %T^rich^ areas in CDC4^WT^ and CDC4^DTR^ tumors at day 14 (N=3 independent experiments; median and interquartile range, Mann-Whitney test). **(c)** Young Modulus of T^rich^ regions and T^poor^ regions in CDC4^WT^ and CDC4^DTR^ tumors at day 14 measured by atomic force microscopy (N=2 independent experiments; median and interquartile range, Mann-Whitney test). **(d)** Image analysis strategy of two-photon, whole slides, SHG/CD3 images. **(e)** Spearman correlation coefficients calculated between fibrillar collagen topography features and the mean number of T cells in 100 µm x 100 µm tiles in CDC4^WT^ and CDC4^DTR^ tumors at day 14 (N=3 independent experiments). **(f)** Principal Component Analysis of local fibrillar collagen topography features in CDC4^WT^ and CDC4^DTR^ tumors at day 14 highlighting: mice genotypes (left) and T^rich^ or T^poor^ classifications (right) (N=3 independent experiments, 7707CC ROIs and 10S70S8 ROIS for CDC4^WT^ and CDC4^DTR^ tumors resp., only 0.1% of ROIs are plotted for representation here). **(g)** Strategy of Random Forest classification of T^rich^ and T^poor^ areas using only collagen topographical features. **(h)** Tables of random forest model performances. **(i)** Permutation analysis showing the importance of energy (60 px and 140 px), coherency (140 px), angle over tumor edge (140 px), and orientation referenced to a 140 px ROI in predicting T cell enrichment in CDC4^WT^ and CDC4^DTR^ datasets (n=10 models; median and interquartile range, Mann-Whitney test). **(j)** Distribution of 140 px ROI SHG mean intensities of CDC4^WT^ and CDC4^DTR^ tumors (N=3 independent experiments, 7707CC ROIs and 10S70S8 ROIS for CDC4^WT^ and CDC4^DTR^ tumors resp.).

**Supplementary figure 3:**
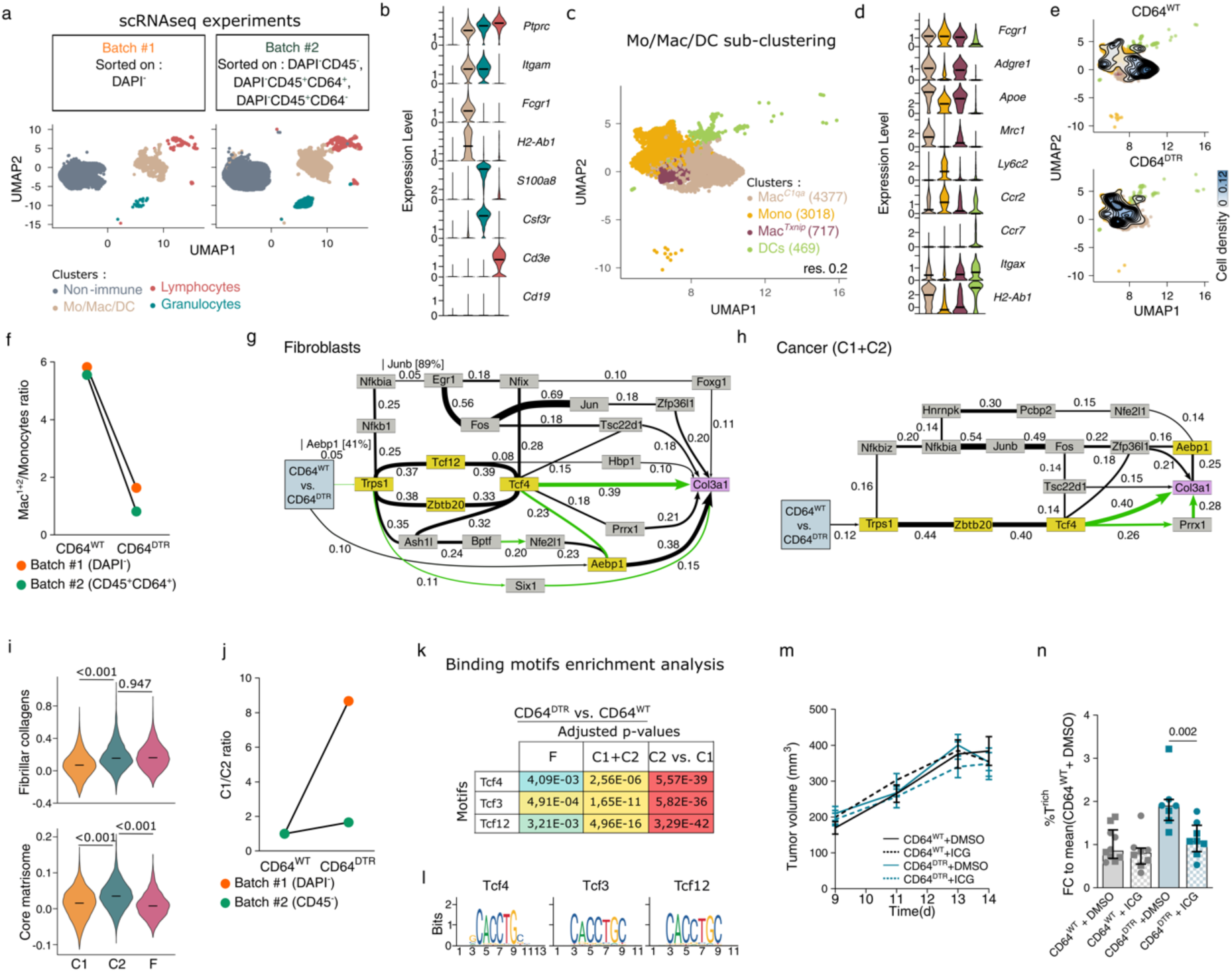
Macrophage depletion induces Col3-fibrotic program driven by Tcf4 in cancer cells and fibroblasts. **(a)** UMAP of merged scRNAseq data from CDC4^WT^ and CDC4^DTR^ tumors from two independent experiments (For batch #1, live cells (DAPI^-^) were sorted by flow cytometry; for batch #2, DAPI^-^CD45^-^, DAPI^-^CD45^+^CDC4^+^, DAPI^-^CD45^+^CDC4^-^ populations were sorted by flow cytometry and sequenced separately). **(b)** Expression levels of classical markers of non-immune cells, monocytes/macrophages/DCs, lymphocytes and granulocytes (line at median). **(c)** UMAP showing sub-clustering of Mo/Mac/DC cluster from CDC4^WT^ and CDC4^DTR^ tumors (merged object from batch #1 and batch #2 DAPI^-^CD45^+^CDC4^+^ samples). 4 clusters were identified by DGEA: Mac^C1qa^, Mono, Mac^Txnip^ and DCs. **(d)** Expression levels of classical markers of monocytes/macrophages/DCs (line at median). **(e)** UMAP of Mo/Mac/DC clusters showing cell densities in CDC4^WT^ and CDC4^DTR^ tumors datasets. **(f)** Ratio between number of macrophages (Mac^C1qa^+Mac^Txnip^) and monocytes in batch #1 and batch #2 DAPI^-^CD45^+^CDC4^-^ samples from CDC4^WT^ and CDC4^DTR^ tumors. **(g)** Transcriptional network induced by macrophage depletion in fibroblasts and **(h)** in cancer cells (C1+C2) identified by MIIC analysis (Top 250 TFs + 44 collagen genes). **(i)** Core matrisome and fibrillar collagen signature expression levels in non-immune clusters (line at median, Wilcoxon test). **(j)** Ratio of Cancer^Airn^ (C2) over Cancer^Lgals3^ (C1) in batch #1 and batch #2 DAPI^-^CD45^-^ samples from CDC4^WT^ and CDC4^DTR^ tumors. **(k)** Tcf4, Tcf3 and Tcf12 adjusted p-values from binding motif enrichment analysis of differentially expressed genes between CDC4^WT^ and CDC4^DTR^ tumors in fibroblasts (F) and cancer cells (C1+C2), and between Cancer^Airn^ (C2) and Cancer^Lgals3^ (C1). **(l)** Logo of Tcf4, Tcf3 and Tcf12 binding motifs from JASPAR database. **(m)** Tumor growth curves from day 9 to day 14 of CDC4^WT^ and CDC4^DTR^ tumors at day 14 treated with Tcf4 inhibitor ICG-001 or DMSO (N=2 independent experiments; mean and SEM, mixed effect analysis). **(n)** %T^rich^ areas (100 µm x 100 µm tiles) in CDC4^WT^ and CDC4^DTR^ tumors at day 14 treated with Tcf4 inhibitor ICG-001 or DMSO (N=2 independent experiments; median and interquartile range, Mann-Whitney test).

**Supplementary figure 4:**
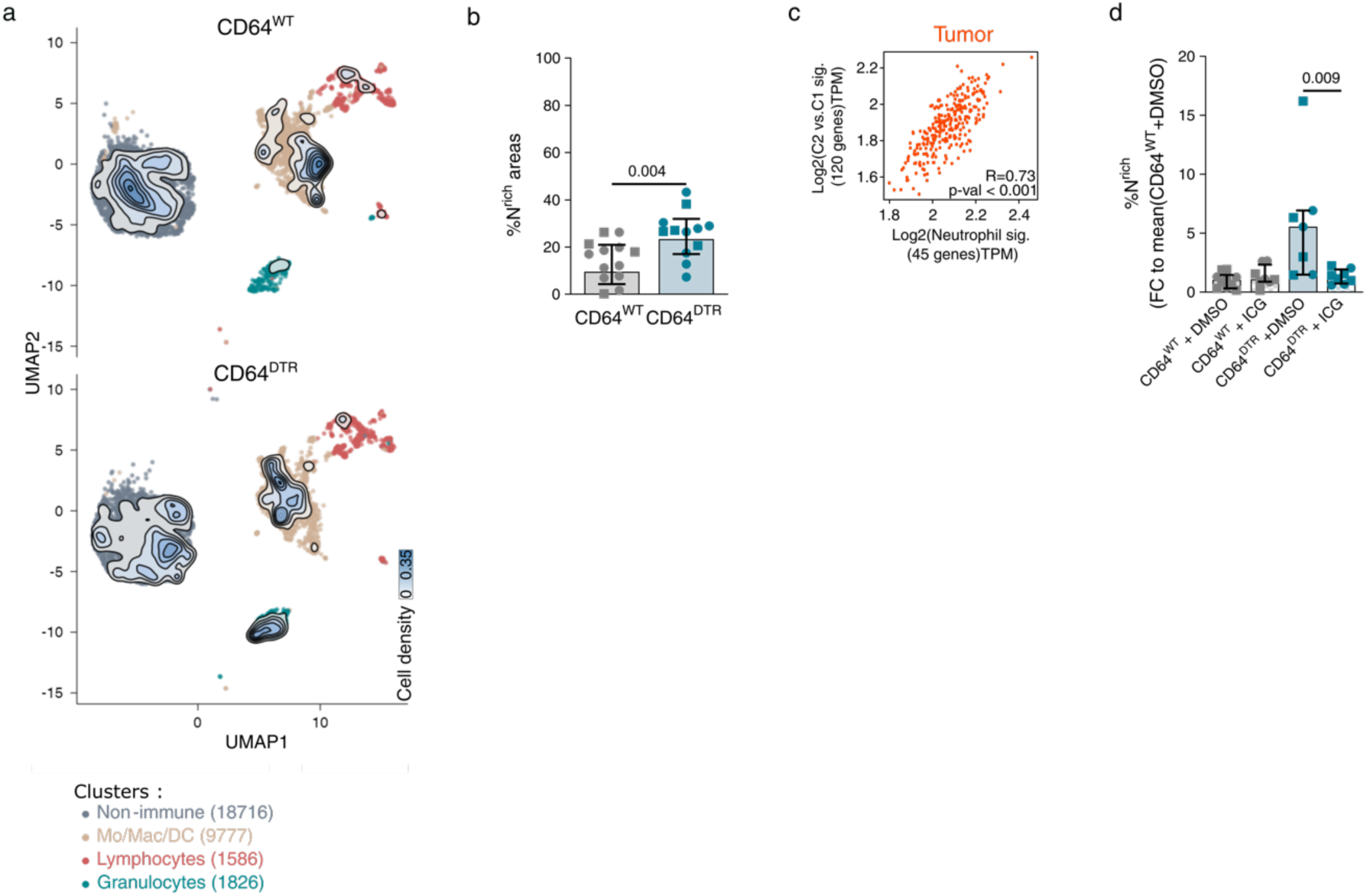
Fibrillar collagen topography dictates tumor overall permissiveness to immune infiltration. **(a)** UMAP of the complete scRNAseq dataset showing cell densities in CDC4^WT^ and CDC4^DTR^ tumors datasets. **(b)** %Neutrophil^rich^ areas (100 µm x 100 µm tiles) in CDC4^WT^ and CDC4^DTR^ tumors at day 14 (N=3 independent experiments; median and interquartile range, Mann-Whitney test). **(c)** Spearman correlation between expression of neutrophil signature and Cancer^Airn^ signature (C2 vs. C1) expression in TCGA COAD tumors bulk RNA sequencing dataset. **(d)** %Neutrophil^rich^ areas (100 µm x 100 µm tiles) in CDC4^WT^ and CDC4^DTR^ tumors at day 14 treated with Tcf4 inhibitor ICG-001 or DMSO normalized to the mean of CD64^WT^ + DMSO condition (N=2 independent experiments; median and interquartile range, Mann-Whitney test).

